# The Nab2 RNA binding protein promotes sex-specific splicing of *Sex lethal* in *Drosophila* neuronal tissue

**DOI:** 10.1101/2020.11.13.382168

**Authors:** Binta Jalloh, J. Christopher Rounds, Brianna E. Brown, Carly L. Lancaster, Sara W. Leung, Ayan Banerjee, Derrick J. Morton, Rick S. Bienkowski, Isaac J. Kremsky, Milo B. Fasken, Anita H. Corbett, Kenneth H. Moberg

## Abstract

The *Drosophila* polyadenosine RNA binding protein Nab2, which is orthologous to a human protein lost in a form of inherited intellectual disability, controls axon projection, locomotion, and memory. Here we define an unexpectedly specific role for Nab2 in regulating splicing of ∼150 exons/introns in the head transcriptome and link the most prominent of these, female retention of a male-specific exon in the sex determination factor *Sex-lethal* (*Sxl*), to a role in m^6^A-dependent mRNA splicing. Genetic evidence indicates that aberrant *Sxl* splicing underlies multiple phenotypes in *Nab2* mutant females. At a molecular level, Nab2 associates with *Sxl* pre-mRNA and ensures proper female-specific splicing by preventing m^6^A hypermethylation by Mettl3 methyltransferase. Consistent with these results, reducing Mettl3 expression rescues developmental, behavioral and neuroanatomical phenotypes in *Nab2* mutants. Overall these data identify Nab2 as a required regulator of m^6^A-regulated *Sxl* splicing and imply a broader link between Nab2 and Mettl3-regulated brain RNAs.

## Introduction

RNA binding proteins (RBPs) play important roles in guiding spatiotemporal patterns of gene expression that distinguish different cell types and tissues within organisms. There are an estimated ∼1500 RBPs that distribute between the nucleus and cytoplasm (Gerstberger et al., 2014), and each has the potential to interact with RNAs to modulate post-transcriptional gene expression. Such regulation is particularly critical in highly specialized cells such as neurons (Conlon and Manley, 2017) where highly regulated alternative splicing of coding regions and 3’UTRs, cleavage/polyadenylation, trafficking and local translation are contribute to precise regulation of gene expression (Brinegar and Cooper, 2016). The critical roles of RBPs in neurons is highlighted by many functional studies that reveal the importance of this class of proteins in brain development and function (Darnell and Richter, 2012) and by the prevalence of human neurological diseases linked to mutations in genes encoding RBPs (Brinegar and Cooper, 2016). Many of these RBPs are ubiquitously expressed and play multiple roles in post-transcriptional regulation. Thus, defining the key neuronal functions of these proteins is critical to understanding both their fundamental roles and the links to disease.

Among the RBPs linked to human diseases are a group of proteins that bind with high affinity to polyadenosine RNAs, which are termed poly(A) RNA binding proteins or Pabs (Kelly et al., 2010). Functional studies of classical nuclear and cytoplasmic Pabs, which utilize RNA recognition motifs (RRMs) to recognize RNA, have uncovered diverse roles for these proteins in modulating mRNA stability, alternative cleavage and polyadenylation and translation (Smith et al., 2014). A second, less well-studied, group of Pabs uses zinc-finger (ZnF) domains to bind target RNAs. Among these is the Zinc Finger Cys-Cys-Cys-His-Type Containing 14 (ZC3H14) protein, which binds with high affinity to poly(A) RNAs via a set of C-terminal tandem Cys-Cys-Cys-His type zinc-finger domains (Leung et al., 2009). ZC3H14 is broadly expressed in many tissues and cell types but mutations in the human *ZC3H14* gene are associated with a heritable form of intellectual disability (Pak et al., 2011), implying an important requirement for this protein in the central nervous system. ZC3H14 has well-conserved homologs in eukaryotes, including *S. cerevisiae* Nuclear poly(A)-binding protein 2 (Nab2), *Drosophila melanogaster* Nab2, *C. elegans* SUT-2 and murine ZC3H14 (Fasken et al., 2019). Zygotic loss of *ZC3H14* in mice and *Drosophila* impairs neuronal function (Pak et al., 2011; Rha et al., 2017), while neuron-specific depletion of *Drosophila* Nab2 is sufficient to replicate these effects (Pak et al., 2011). Reciprocally, expression of human ZC3H14 in Nab2-deficient neurons rescues this defect, demonstrating a high degree of functional conservation between human ZC3H14 and *Drosophila* Nab2 (Kelly et al., 2014). Collectively, these data focus attention on what are critical, but poorly understood, molecular roles for ZC3H14/Nab2 proteins in neurons.

Neuronal ZC3H14/Nab2 can be divided into two pools, a nuclear pool that accounts for the majority of ZC3H14/Nab2 in the cell, and a small cytoplasmic pool of protein detected in mRNA ribonucleoprotein particles (mRNPs) of axons and dendrites (Bienkowski et al., 2017; Leung et al., 2009; Rha et al., 2017). Depletion of both pools in *Drosophila* neurons cause defects in axon genesis within the brain mushroom bodies (Kelly et al., 2016), a pair of twin neuropil structures involved in learning and memory (Armstrong et al., 1998; Heisenberg, 2003). This requirement has been linked to a physical association between cytoplasmic Nab2 and the *Drosophila* Fragile-X mental retardation protein homolog, Fmr1 (Wan et al., 2000), and translational repression of shared Nab2-Fmr1 target RNAs in the cytoplasm (Bienkowski et al., 2017). Despite this insight into a cytoplasmic function of Nab2, molecular roles of the abundant population of Nab2 in neuronal nuclei remain undefined.

Here, we employed a broad and an unbiased RNA sequencing approach to identify transcriptome-wide changes in the heads of *Nab2* loss-of-function mutant flies. While the steady-state levels of most transcripts were not significantly changed, we uncovered a striking effect on splicing of a subset of neuronal RNA transcripts. We focused our analysis on a well-characterized sex-specific alternative splicing event in the *Sex-lethal* (*Sxl*) transcript. Results reveal that *Nab2* plays a novel role in regulating the alternative splicing of *Sxl* in a sex-specific manner. Recent works has revealed a role for m^6^A RNA methylation by the enzyme Mettl3 in modulating this splicing event (Kan et al., 2017; Lence et al., 2016). Similar to *Mettl3*, the requirement for *Nab2* in alternative splicing of *Sxl* is only essential for neuronally-enriched tissues. Genetic and biochemical experiments support a functional link between m^6^A methylation and Nab2 function. These results demonstrate the role for *Drosophila Nab2* in RNA alternative splicing as well as RNA methylation and sex determination in the neurons.

## Results

### Nab2 loss affects levels and processing of a subset of RNAs in the head transcriptome

To assess the role of Nab2 in regulating the central nervous system transcriptome, a high-throughput RNA Sequencing (RNA-Seq) analysis was carried out in triplicate on *Nab2* null mutant heads (*Nab2^ex3^* imprecise excision of *EP3716*) (Pak et al., 2011) and isogenic control heads (*Nab2^pex41^* precise excision of *EP3716*). To capture sex-specific differences, heads were collected from both male and female flies of each genotype. Briefly, total RNA from 1-day old adults was rRNA-depleted and used to generate stranded cDNA libraries that were sequenced (150 cycles) on a NextSeq 500 High Output Flow Cell. This generated a total of approximately 1.1 billion 75 base-pair (bp) paired-end reads (91 million/sample) that were mapped onto the Dmel6.17 release of the *Drosophila* genome using RNA STAR (Dobin et al., 2013). Read annotation and per-gene tabulation was conducted with featureCounts (Liao et al., 2014) and differential expression analysis was performed with DESeq2 (Love et al., 2014).

RNA sequencing reads across the *Nab2* gene are almost completely eliminated in *Nab2^ex3^* mutants, confirming the genetic background and integrity of the analysis pipeline (**Supplemental Figure 1**). Principal component analysis (PCA) performed with DESeq2 output data confirms that the 12 RNA-seq datasets distribute into four clusters that diverge significantly from one another based on genotype (*Nab2^ex3^* vs. *Nab2^pex41^* control; PC1 58% variance) and sex (male vs. female; PC2 26% variance) (**Figure 1A**). The DESeq2 analysis detects 3,799 and 1,545 genes in females and males, respectively, that exhibit statistically significant differences in RNA abundance between *Nab2^ex3^* and control (Benjamini-Hochberg [BH] adjusted *p*-value/false discovery rate (FDR)<0.05). Comparison of fold-changes (*Nab2^ex3^* vs. control) among these significantly different RNAs reveals a high degree of correlation in female vs. male samples (R=0.79), particularly among RNAs whose levels are most elevated upon Nab2 loss (**Figure 1B**). Applying a 2-fold change cutoff (|log_2_[fold-change]|*≥*1) trims these sets to 453 significantly changed RNAs in females (294 ‘up’, 159 ‘down’), and 305 significantly changed RNAs in males (150 ‘up’, 155 ‘down’) (**Figure 1C**), which merge into a combined set of 570 significantly affected RNAs that trend similarly in heatmap analysis of mutant vs. control samples (**Figure 1D**). A majority of the 453 affected ‘female’ RNAs are mRNAs (439) and the remaining are snoRNAs (8), snRNAs (1), pre-rRNAs (1), and tRNAs (4) (**Figure 1E**). A similar distribution occurs in male heads: a majority of the affected RNAs are mRNAs (297) and the remainder are snoRNAs (4), snRNAs (1), pre-rRNAs (1), and tRNAs (2) (**Figure 1E**). Overall, the number of significantly changed RNAs ((|log_2_[fold-change]|*≥*1 and FDR<0.05) in *Nab2^ex3^* females and male heads represents a small fraction of RNAs detected in heads (2.2% and 3.7% in males and females, respectively), suggesting that Nab2 normally contributes to RNA-specific regulatory mechanisms in *Drosophila* head tissue.

**Figure 1.**
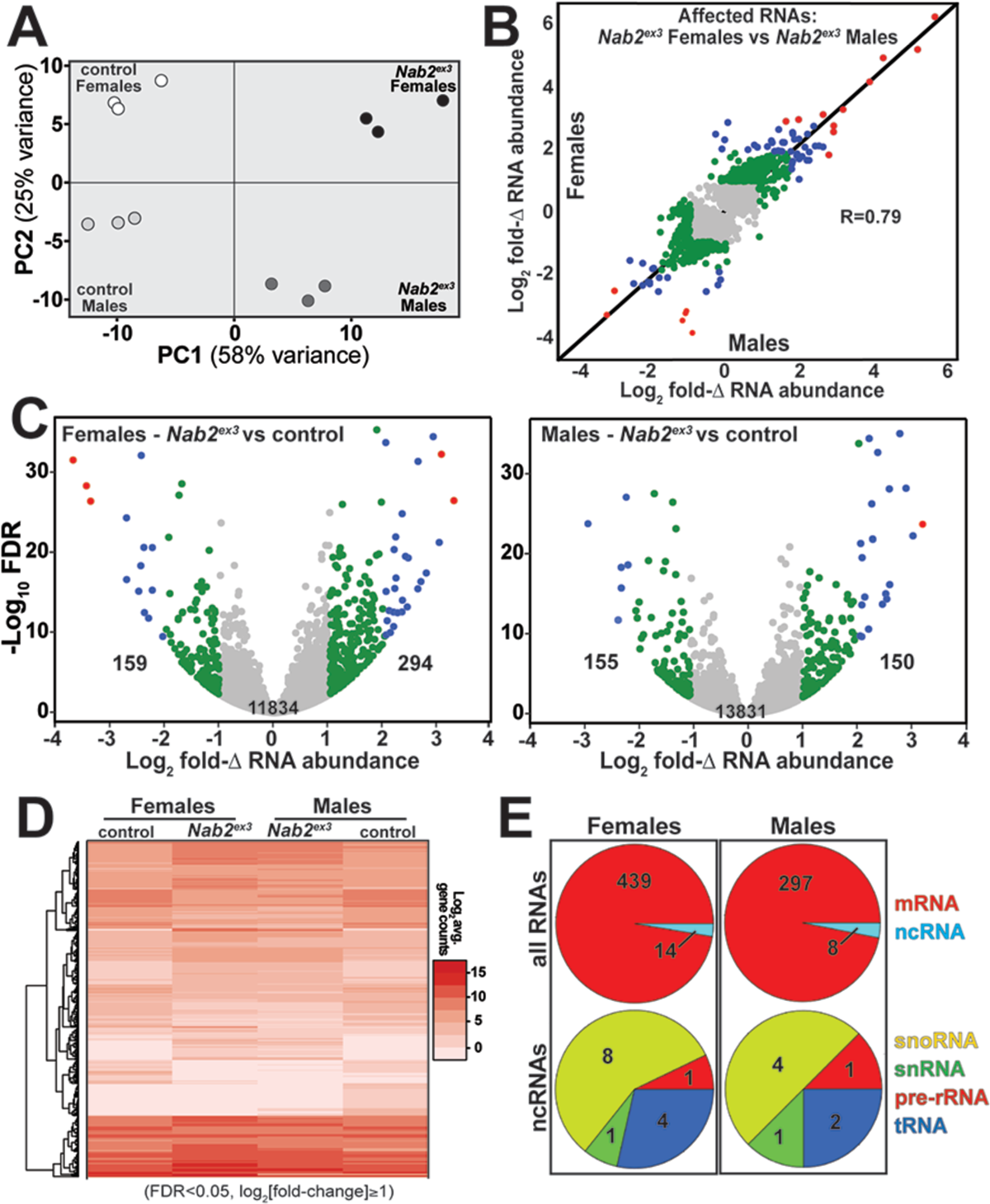
RNA sequencing detects effects of Nab2 loss on the head transcriptome. **(A)** Principal component analysis (PCA) of RNA-seq data from three biological replicates of control and *Nab2* mutant (*Nab2^ex3^*) male and female heads. **(B)** Correlation scatter plot of log_2_ fold-change (Δ) in abundance of affected RNAs in males and females (log_2_ average gene counts: **grey**<1, 1≤**green**<2, 2≤**blue**<3, **red**≥3). **(C)** Volcano plots of fold-Δ in abundance vs. false discovery rate (FDR -log_10_) of affected RNAs in *Nab2^ex3^* females and males (dot plot color coding as in B). Elevated (≥1), reduced (≤-1), and total RNAs are indicated. **(D)** Heatmap comparison of significantly changed gene counts (FDR<0.05; |log_2_ fold-Δ|≥1) in *Nab2^ex3^* females and males vs. sex-matched controls. **(E)** Pie chart shows distribution of RNA classes among significantly affected RNAs detected in **C** and **D**.

### Nab2 loss alters levels of transcripts linked to mRNA processing

To screen Nab2-regulated RNAs for enriched functions, Gene Set Enrichment Analysis (GSEA) (Mootha et al., 2003; Subramanian et al., 2005) was carried to identify enriched gene ontology (GO) terms (Ashburner et al., 2000; The Gene Ontology, 2019) among the significantly changed female and male RNAs identified by DESeq2. This filtering uncovers significant enrichment (*p*<0.05) for “RNA splicing” GO (GO:0008380) within the upregulated group of RNAs in both sexes (**Figure 2A**). In *Nab2^ex3^* females, 32 of 155 genes annotated under this term are present among upregulated RNAs; whereas in males, 75 of 159 genes annotated under this term are present among upregulated RNAs (**Figure 2A**). This enrichment for upregulated splicing-related factors indicates that Nab2 loss could shift splicing patterns in the adult head. Consistent with this hypothesis, MISO (mixture of isoforms) analysis (Katz et al., 2010) of annotated alternative splicing events confirms that Nab2 loss significantly alters splicing patterns within a small number of transcripts in female (50) and male (51) heads (**Supplemental Table 1**) that fall into a variety of GO terms (**Supplemental Figure 2**). These MISO-called alternative splicing events include 5’ and 3’ alternative-splice site usage, intron retention events, and previously annotated exon skipping events, some of which are detected in the same transcripts (**Figure 2B**).

**Figure 2.**
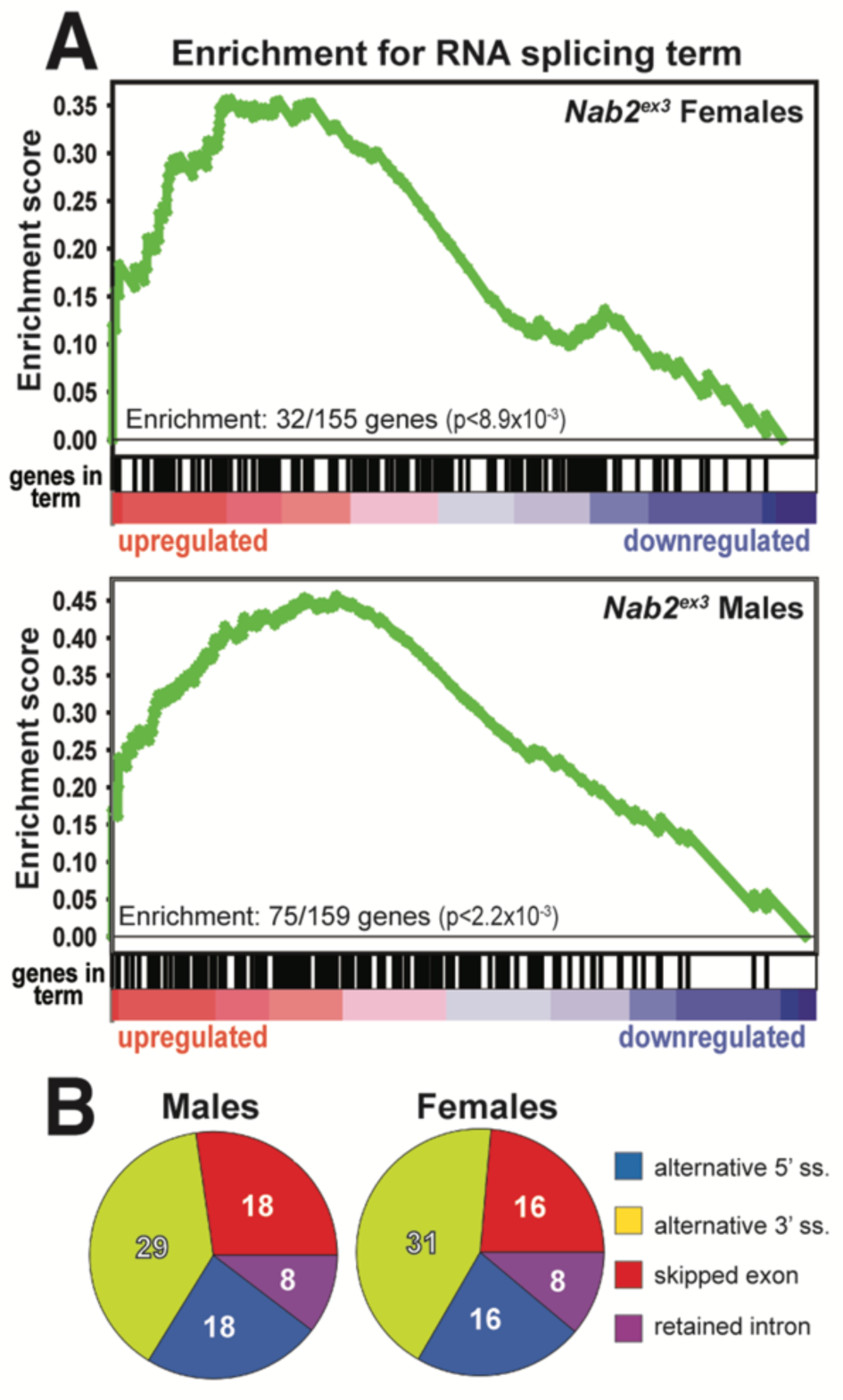
Significantly up/down-regulated RNAs in *Nab2^ex3^* heads are enriched for predicted splicing factors. **(A)** Gene set enrichment analysis (GSEA) detects enrichment for the ‘RNA splicing’ GO term in up- and down-regulated gene sets in both female (top) and male (bottom) *Nab2^ex3^* datasets. Gene enrichments are indicated with corresponding p-values. **(B)** Pie chart illustrating the distribution of previously annotated alternative splicing RNA splicing events that are significantly altered in *Nab2^ex3^* mutant female and male heads (ss=splice site).

To test whether Nab2 loss results in unannotated or aberrant splicing events, DEXSeq analysis (Anders et al., 2012) was performed to scan for differential abundance of individual exons relative to other exons within the same transcript. This analysis detects 151 affected RNAs in *Nab2^ex3^* females and 114 in *Nab2^ex3^* males (**Table 1**), with many top-ranked transcripts encoding factors with roles in behavior, neurodevelopment, and/or neural function (**Supplemental Table 2**).

**Table 1.**
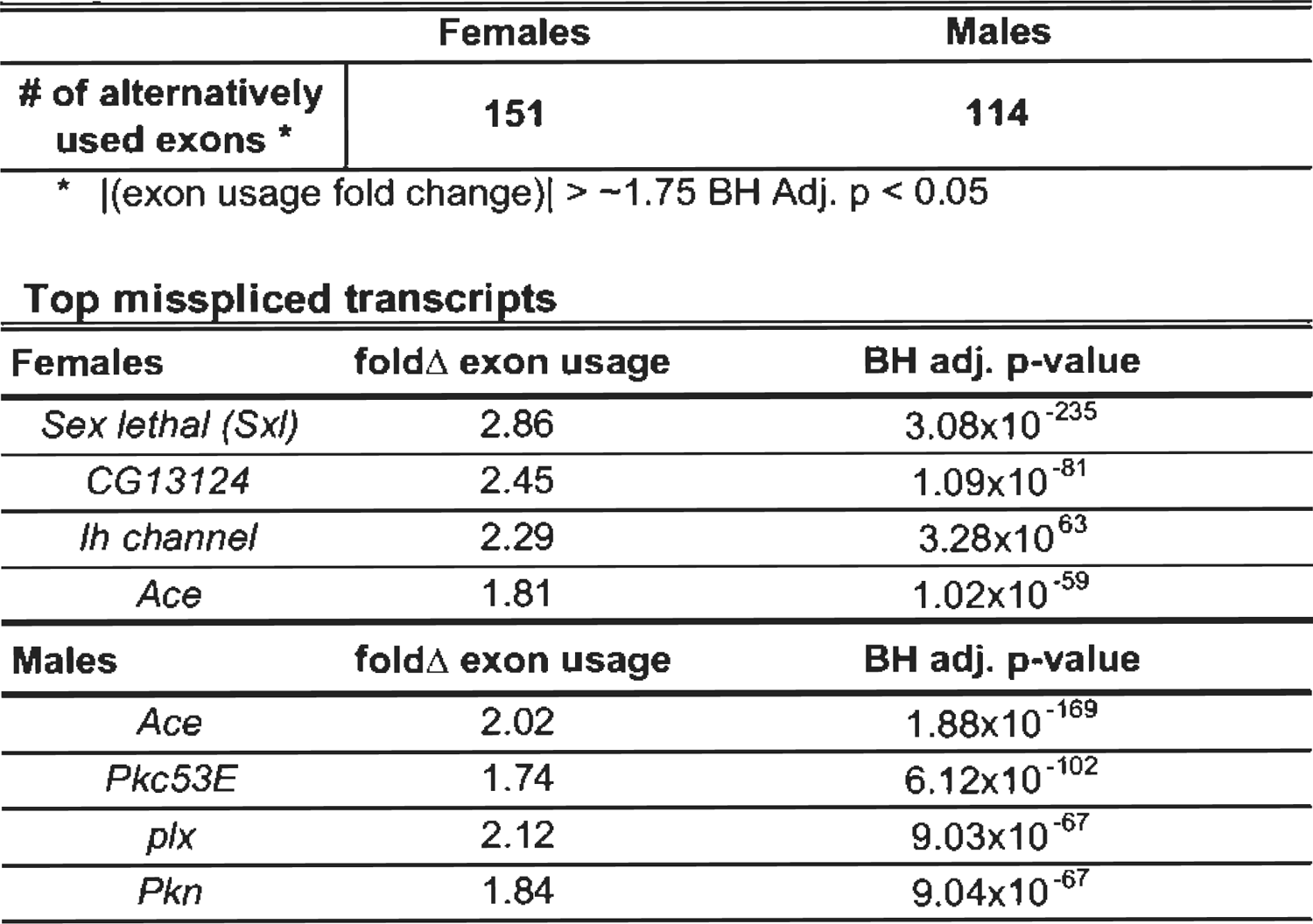
Alternative exon usage (DEXSeq) in *Nab2*^ex3^ head transcriptomes

The most statistically significant exon usage change in either sex is female-specific inclusion of exon 3 in the *Sex lethal* (*Sxl*) mRNA (2.86-fold increase, *p*=3.08×10^-235^). This effect on *Sxl* mRNA in *Nab2^ex3^* females is followed in rank order of significance by enhanced inclusion of exons 1 and 2 of the MIF4GD homolog transcript *CG13124*, exons 1 and 2 of the voltage-gated ion channel transcript *I_h_ channel* (*I_h_*), and exon 1 of the synaptic enzyme transcript *Acetylcholine esterase* (*Ace)*. In *Nab2^ex3^* males, the top four events are enhanced inclusion of exon 1 of the *Ace* transcript, exon 1 of the *Protein kinase C at 53E* (*Pkc53E*) transcript, exons 1 and 2 of the Rab GTPase *pollux* (*plx*) transcript, and exons 1 and 2 of *Protein kinase N* (*Pkn*) transcript. In some cases, identical exons are affected in both *Nab2^ex3^* sexes and accompanied by retention of the intervening intron (e.g. see *CG13124* and *I_h_* traces in **Supplemental Figure 2**). The robust increase in *Sxl* exon 3 in *Nab2^ex3^* females is noteworthy both for the central role that differential inclusion of exon 3 plays in *Drosophila* sex determination (Harrison, 2007), but also because DEXSeq did not detect changes in exon 3 inclusion or abundance in *Nab2^ex3^* males. In light of this sex-specific effect of Nab2 loss on alternative splicing of *Sxl* exon 3, subsequent analyses focused on the role of Nab2 in *Sxl* mRNA splicing in female heads.

### *Nab2^ex3^* females exhibit masculinized *Sxl* splicing in neuron-enriched tissues

The Sex lethal (Sxl) protein is a female-specific, U-rich RNA binding protein that acts through the *tra-dsx* and *msl-2* pathways to promote female somatic and germline identity (Gawande et al., 2006; Penalva and Sanchez, 2003). *Sxl* pre-mRNA is expressed in both males and females, but alternative splicing regulated by m^6^A RNA methylation and several RBPs leads to female-specific skipping of exon 3 during splicing (Haussmann et al., 2016; Lence et al., 2016; Sakamoto et al., 1992). Because exon 3 includes an in-frame translation ‘stop’ codon, full-length Sxl protein is only made and active in female cells (Bell et al., 1988). The inclusion of *Sxl* exon 3 in *Nab2^ex3^* mutants would thus implicate Nab2 as a novel component of molecular machinery that controls *Sxl* pre-mRNA splicing in female heads.

Visualizing *Sxl* RNA-Seq reads with IGV Viewer (Robinson et al., 2017) confirms a large increase in exon 3 reads in *Nab2^ex3^* females (*Nab2^ex^*^3^-F) relative to control females (control-F), and also shows retention of ∼500 bases of intron 3 sequence in *Nab2^ex3^* females (**Figure 3A**). Quantification of reads across the entire *Sxl* locus detects an ∼1.5-fold increase in the overall abundance of the *Sxl* mRNA in *Nab2^ex^*^3^ females compared to control females. Normal splicing patterns are detected across all other *Sxl* intron-exon junctions in both genotypes of males and females, including female-specific exon 9 inclusion (**Figure 3A**). Reverse transcription polymerase chain reaction (RT-PCR) on fly heads using *Sxl* primers (see arrows in **Figure 3A** schematic) that detect exon 2-exon 4 (control females) and exon 2-exon 3-exon 4 (control males) confirms the presence of the mis-spliced exon 2-exon 3-exon 4 mRNA transcript in *Nab2^ex3^* females (**Figure 3B**). The exon 2-exon 3-exon 4 mRNA transcript appears to be more abundant in *Nab2^ex3^* female heads than in female heads lacking *Mettl3*, which encodes the catalytic component of an m^6^A methyltransferase complex that promotes exon 3 skipping in nervous system tissue (Haussmann et al., 2016; Kan et al., 2017; Lence et al., 2016). RT-PCR also reveals a ∼1kb band in *Nab2^ex3^* females (arrowhead, **Figure 3B**) that sequencing identifies as aberrantly spliced transcript that incorporates 503 bases of intron 3 leading up to a cryptic 5’ splice site (i.e. exon 2-exon 3-intron 3^503^-exon 4); this matches the *Sxl* intron 3 sequencing reads observed in IGV (see **Figure 3A**). Significantly, RT-PCR analysis of *Sxl* mRNA in dissected control and *Nab2^ex3^* females detects exon 3 retention in *Nab2^ex3^* thoraxes, but not in abdomens or ovaries (**Figure 3C**). This result implies that Nab2 is only necessary to direct *Sxl* exon 3 exclusion in specific tissues or cell types such as neurons, which are enriched in the head (brain) and thorax (ventral nerve cord). In sum, these data reveal a tissue-specific role for Nab2 in blocking *Sxl* exon 3 inclusion in females and regulating 5’-splice site utilization at the exon 3-exon 4 junction.

**Figure 3.**
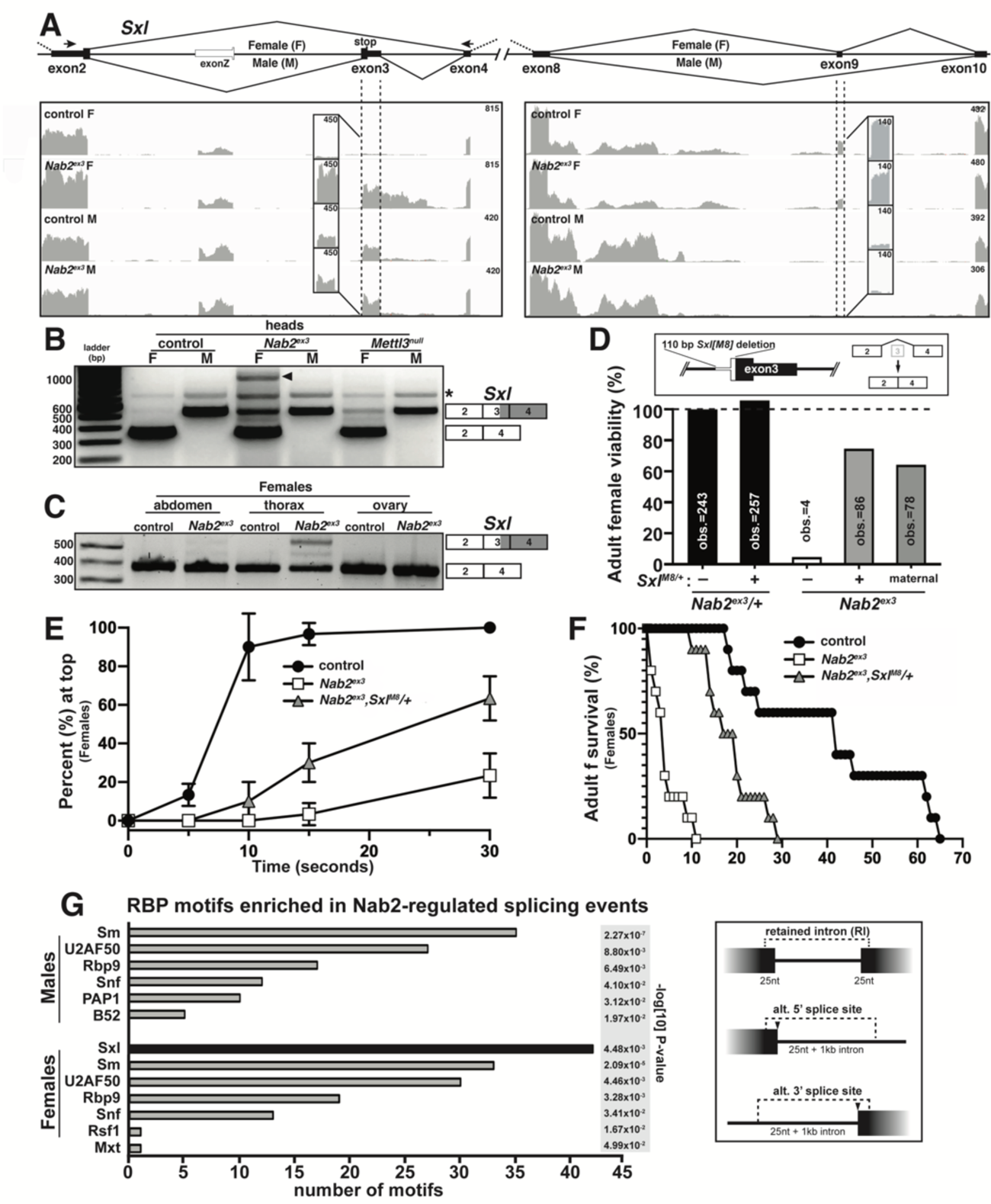
Alternative splicing of *Sxl* is disrupted in *Nab2^ex3^* female heads. **(A) Top panel**: normal *Sxl* alternative splicing patterns across the exon 2-4 and exon 8-10 regions in females (F) and males (M). Arrows indicate location of primers used in **B**. **Bottom panel**: corresponding sequencing reads across the *Sxl* locus in the indicated sexes and genotypes. Dotted lines and boxed insets highlight exon 3 and exon 9 reads. **(B)** RT-PCR analysis of *Sxl* mRNA in control, *Nab2^ex3^* and *Mettl3^null^* male (M) and female (F) heads. Exon 2-3-4 and exon 2-4 bands are indicated. Arrowhead denotes exon 2-3-intron-4 product noted in text. Asterisk (*) is non-specific product. **(C)** RT-PCR analysis of *Sxl* mRNA in adult female (F) control and *Nab2^ex3^* tissues using the same primers as in **B** and with exon 2-3-4 and 2-4 bands indicated. **(D)** A single copy of the *Sxl^M8^* allele, which harbors a 110 bp deletion that causes constitutive exon2-4 splicing, partially suppresses lethality of Nab2^ex3^ females, both zygotically and maternally. **(E-F)** *Sxl^M8^* dominantly (i.e. *M8/+*) suppresses previously defined locomotion (as assessed by negative-geotaxis) and life-span defects among age-matched *Nab2^e3^* females. **(G)** RNA binding protein (RBM) motif enrichment analysis detects predicted Sxl binding sites as the most frequent motif among Nab2-regulated splicing events in female heads. Other enriched motifs are similar between male and female heads. Regions used for motif analysis (retained introns, and alternative 5’ or 3’ splice sites plus flanking sequence) are shown in the schematic.

*Sxl* exon 3 inclusion in *Nab2^ex3^* female head RNAs suggests that insufficient levels of the exon 2-exon 4 splice variant contribute to *Nab2^ex3^* phenotypes in females. To test this hypothesis, the constitutively female-spliced *Sxl^M8^* allele (Barbash and Cline, 1995) was placed as a heterozygote into the background of *Nab2^ex3^* animals. *Sxl^M8^* contains a 110 bp deletion covering the 5’-end of intron 2 and 3’-end of exon 3 and consequently undergoes constitutive splicing to the feminized exon 2-exon 4 variant regardless of sex (top panel, **Figure 3D**). Heterozygosity for this *Sxl^M8^* allele produces strong rescue of *Nab2^ex3^* mutant female viability from ∼4% to 71% (*Sxl^M8^/+;;Nab2^ex3^*) (**Figure 3D**). Female *Nab2^ex3^* siblings that did not inherit the *Sxl^M8^* allele also exhibit elevated viability (64%), perhaps due to maternal loading of *Sxl* mRNA (**Figure 3D**). Surviving *Sxl^M8^/+;;Nab2^ex3^* females also show improved locomotion in a negative geotaxis assay (**Figure 3E**) and lengthened lifespan (**Figure 3F**) relative to *Nab2^ex3^* females. Consistent with the original report describing *Sxl^M8^* (Barbash and Cline, 1995), the allele is male-lethal in control (*Nab2^pex41^*) and *Nab2^ex3^* backgrounds. This female-specific rescue of *Nab2^ex3^* by *Sxl^M8^* indicates that restoring Sxl expression can compensate for Nab2 loss in some developing tissues As Sxl is itself an RBP with roles in alternative splicing (Bell et al., 1988; Penalva and Sanchez, 2003), the rescuing effect of the *Sxl^M8^* allele prompted a bioinformatic scan for RBP motifs enriched in proximity to the Nab2-dependent alternative splicing events identified by MISO analysis (see **Figure 2B**). Input sequences were composed of retained introns plus 25bp extending into each flanking exon, and alternative splice sites with 25bp of exon plus 1kb of adjacent intron (see schematic, **Figure 3G**). This unbiased scan detected predicted Sxl binding sites as the single most abundant RBP motif within the Nab2-regulated MISO events in females (**Figure 3G**). Notably, Sxl motifs were not detected as enriched in the male *Nab2^ex3^* MISO dataset, which otherwise strongly resembles the remaining group of female-enriched RBP motifs (e.g. the HNRNPL homolog *smooth* (*sm*), *RNA binding protein-9* (*Rbp9*), the U1-SNRNPA homolog *sans fille* (*snf*), and the U2-SNRNP component *U2AF50*). The female-specific enrichment for Sxl binding sites suggests that Nab2 may regulate alternative splicing events indirectly via control of a *Sxl*-regulated splicing program. Intriguingly, the Sxl target *transformer* (*tra*) and the Tra target *double-sex* (*dsx*) (Horabin and Schedl, 1993; Sanchez et al., 2001) were not recovered in the *Nab2^ex3^* MISO or DESeq2 datasets, and IGV reads show no evidence of altered structure of their RNAs relative to *Nab2^pex41^* controls (**Supplemental Figure 4**). Together these data suggest that Sxl may not control the *tra*-*dsx* pathway in the adult head, or that *tra* and *dsx* splicing are only altered in a subset of *Nab2^ex3^* head cells and thus not detectable by bulk RNA-Seq analysis.

### The dosage compensation complex contributes to phenotypes in *Nab2^ex3^* mutant females

The lack of evidence that *tra* and *dsx* mRNAs are affected by *Nab2* loss prompted analysis of the other major role of Sxl, which is to bind to the *male-specific lethal-2* (*msl-2*) mRNA and inhibit its translation in female somatic and germline tissues (Keller and Akhtar, 2015; Lucchesi and Kuroda, 2015). As a result, Msl-2 protein is only expressed in male cells, where it promotes assembly of a chromatin modifying complex termed the <u>D</u>osage <u>C</u>ompensation <u>C</u>omplex (DCC; composed of Msl-1, Msl-2, Msl-3, Mof, Mle and *roX1* and *roX2* non-coding RNAs), which is recruited to the male X chromosome to equalize X-linked gene expression between males and females (Keller and Akhtar, 2015; Lucchesi and Kuroda, 2015). A number of DCC components are expressed highly in the adult nervous system (Amrein and Axel, 1997), which correlates with the tissue-restricted link between Nab2 and *Sxl* splicing (as in **Figure 3B**). As a functional test of interactions between Nab2 and the DCC pathway, a loss-of-function allele of *msl-2* (*msl-2^killer of males-A^* or *msl-2^kmA^*) (Bevan, 1993) was tested for dominant effects on *Nab2^ex3^* female phenotypes. Remarkably, a single copy of *msl-2^kmA^* significantly rescues defects in viability (**Figure 4A**), lifespan (**Figure 4B**), and locomotion (**Figure 4C**) among *Nab2^ex3^* females. *roX1* and *mle* loss-of-function alleles were also able to rescue *Nab2^ex3^* phenotypes (**Supplemental Figure 5**). Given that Msl-2 is not normally active in adult female tissues (Amrein and Axel, 1997; Meller et al., 1997) and that forced *msl*-2 expression reduces female viability (Kelley et al., 1995), rescue by *msl-2^kmA^* heterozygosity provides strong evidence that the DCC pathway is inappropriately activated in *Nab2^ex3^* females. Of note, the *msl-2* and *mle* RNAs appear similar in IGV reads from control and *Nab2^ex3^* adults (**Supplemental Figure 4**), indicating that genetic interactions between these loci are not likely due to direct effects of Nab2 loss on abundance and structure of these RNAs.

**Figure 4.**
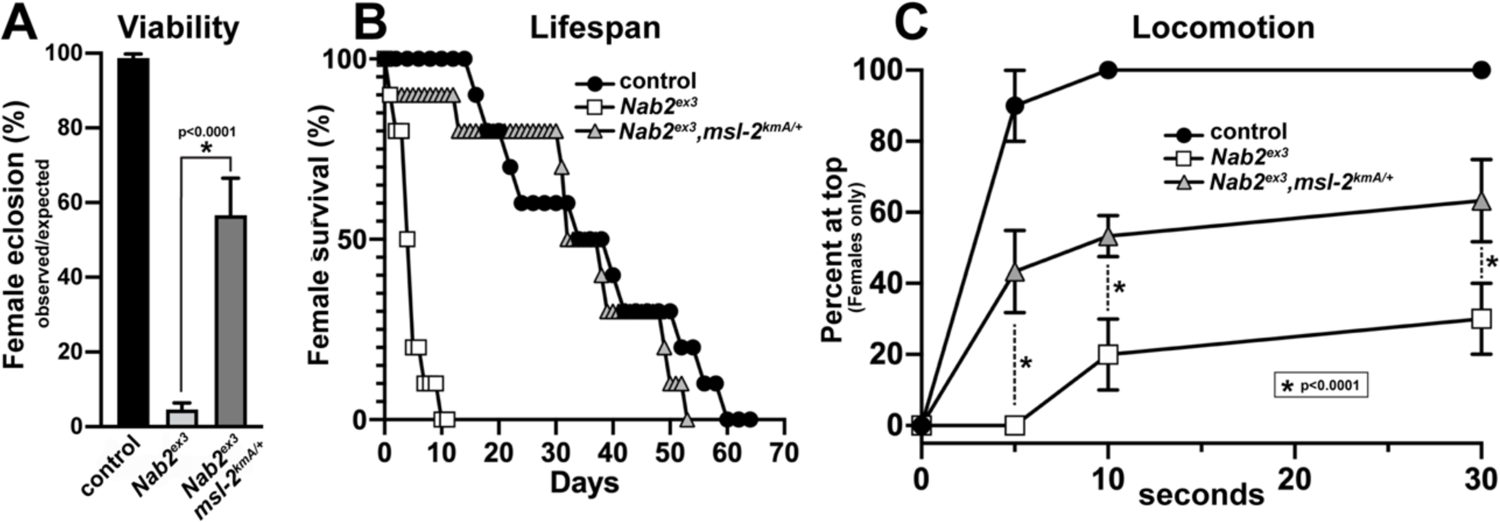
An allele of the DCC component *male-specific lethal-2* (*msl*-2) rescues *Nab2* phenotypes in females. **(A)** Percent of control, *Nab2^ex3^*, and *msl-2^kmA^/+;;Nab2^ex3^* (*msl*-2 is on the X chromosome) females eclosing as viable adults (calculated as #observed/#expected). **(B)** Survival of age-matched adult female flies of the same genotypes. **(C)** Negative geotaxis of age-matched adult females of the same genotypes at 5sec, 10sec and 30 sec timepoints. Significance values are indicated (\**p*<0.0001).

### Nab2-regulated splicing of *Sxl* exon3 is dependent upon the Mettl3 m^6^A methyltransferase

Genetic interactions between *Nab2*, *Sxl*, and *msl-2* alleles are consistent with a role for Nab2 protein in regulating sex-specific splicing of *Sxl* exon 3. One mechanism that promotes exon 3 exclusion in females is based on N^6^-methylation of adenosines (m^6^A) in *Sxl* pre-mRNA by the Methyltransferase like-3 (Mettl3)-containing methyltransferase complex (reviewed in Lence et al., 2017). Inactivating mutations in components of this m^6^A ‘writer’ complex masculinize the pattern of exon 3 splicing in female flies (Haussmann et al., 2016; Kan et al., 2017; Lence et al., 2016) in a manner similar to *Nab2^ex3^*. Molecular studies indicate that the Mettl3 complex promotes exon 3 exclusion in females by depositing m^6^A within *Sxl* exon 3 and flanking introns (Haussmann et al., 2016; Kan et al., 2017; Lence et al., 2016).

To assess Nab2-Mettl3 functional interactions, the null allele *Mettl3^null^* (formerly known as *Ime4^null^)* (Lence et al., 2016) was recombined into the *Nab2^ex3^* background (**Supplemental Figure 6**; the loci are 281kb apart on chr3R), and then used to test for effects on phenotypes in *Nab2^ex3^,Mettl3^null^* double mutant females. As described previously (Haussmann et al., 2016; Kan et al., 2017; Lence et al., 2016), loss of Mettl3 in an otherwise *wildtype* genetic background reduces adult viability, shortens lifespan and decreases locomotion in a negative geotaxis assay (**Figure 5A-C**). However, removing *Mettl3* has the inverse effect of suppressing each of these defects in *Nab2^ex3^* females (**Figure 5A-C**): *Nab2^ex3^,Mettl3^null^* double mutant females show approximately 5-fold higher viability, 1.5-fold longer lifespan, and 2-fold greater locomotion activity (at the 30sec time point; **Figure 5C**) than *Nab2^ex3^* mutants alone. Significantly, qPCR analysis confirms that *Mettl3* loss causes aberrant *Sxl* exon 3-exon 4 splicing (as reported in Haussmann et al., 2016; Kan et al., 2017; Lence et al., 2016) but reduces it in *Nab2^ex3^,Mettl3^null^* double mutant females relative to *Nab2^ex3^* females (**Figure 5D-E**). Within the adult brain, removing *Mettl3* also rescues structural defects in *α*- and *β*-lobes in the mushroom bodies (MBs) that are otherwise highly penetrant in *Nab2^ex3^* adults (**Figure 5F**). *Nab2^ex3^* brains normally show 60-80% penetrance of thinned or missing *α*-lobes and fused *β*-lobes (Bienkowski et al., 2017; Kelly et al., 2016), but *Nab2^ex3^,Mettl3^null^* double mutants exhibit a reduction of *α*-lobe defects and complete suppression of the *β*-lobe fusion defect (**Figure 5F**). As Nab2 and Mettl3 each act autonomously within MB neurons to pattern *α*/*β*-lobe structure (Kelly et al., 2016; Soldano et al., 2020), these genetic interactions imply that Nab2 and Mettl3 may co-regulate pathways which guide axon projection.

**Figure 5.**
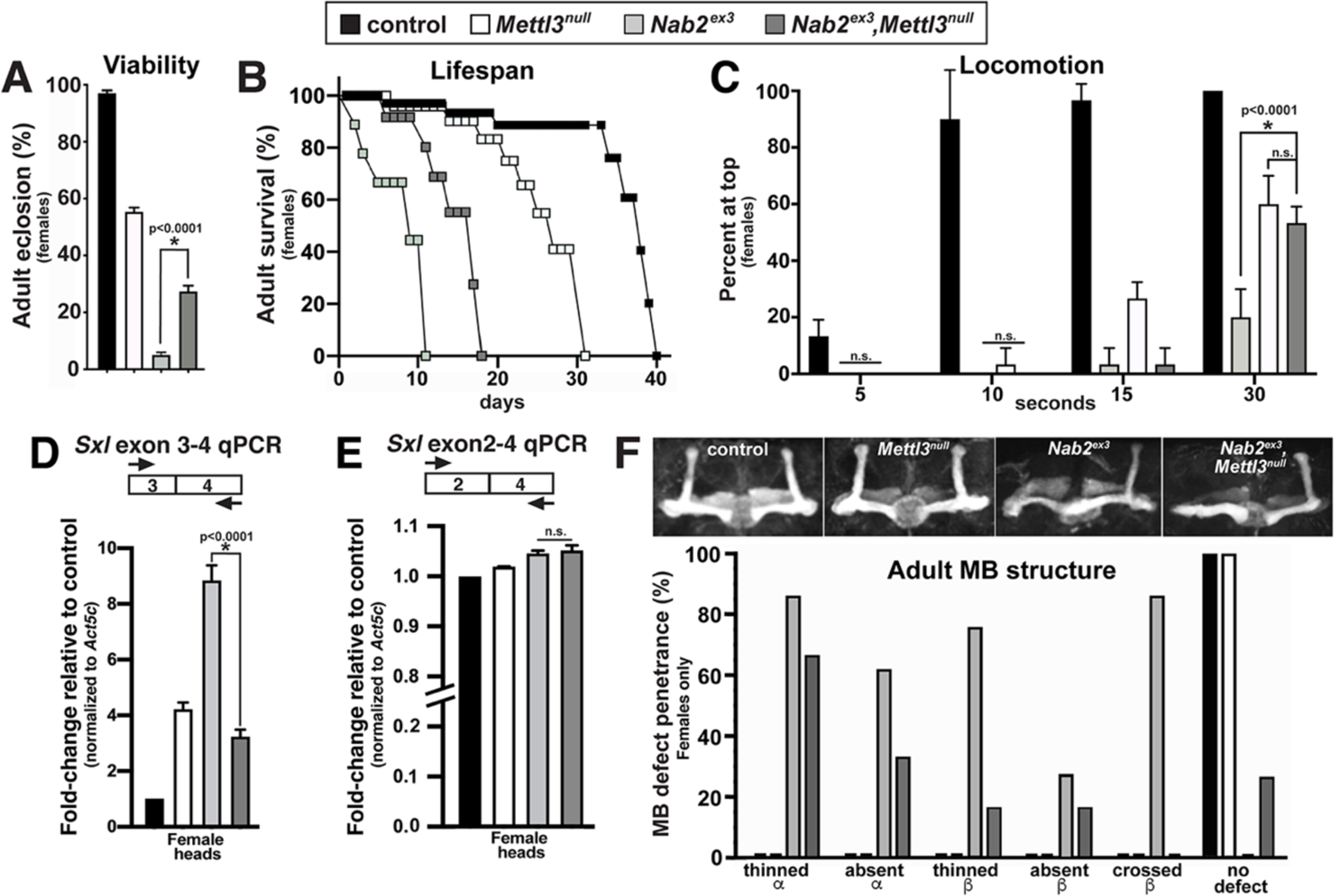
Removing the Mettl3 m^6^A transferase suppresses viability, behavioral, neuroanatomical and *Sxl* splicing defects in *Nab2* mutant females. Color coding as indicated: control (**black** fill), *Mettl3^null/null^* ( fill), *Nab2^ex3/3×3^* fill), and *Nab2^ex3/ex3^,Mettl3^null/null^* (fill). **(A)** Percent of control, *Nab2^ex3^* and *Nab2^ex3^,Mettl3^null^* females eclosing as viable adults (calculated as #observed/#expected). **(B)** Survival of age-matched adult female flies of the indicated genotypes. **(C)** Negative geotaxis of adult females of the indicated genotypes at indicated timepoints. Significance values are indicated at the 30sec timepoint (* *p*<0.0001; n.s.=not significant). **(D-E)** Quantitative real-time PCR analysis of the abundance of *Sxl* mRNA variants containing **(D)** the E3-E4 (exon3-exon4) or **(E)** the E2-E4 splice forms in female heads of the indicated genotypes. Primers denoted by arrows in the cartoon schematic. CT values are relative to control (*Nab2^pex41^*) and normalized to *Act5c* control (\**p*<0.0001; n.s.=not significant). **(F) Top panel:** representative Z-stacked confocal images of anti-Fas2 staining to visualize MBs in female brains of indicated genotypes. **Bottom panel:** penetrance of MB phenotypes (thinned/absent *α*-lobes, thinned/absent *β*-lobes, and *β*-lobes mid-line crossing) in adult females of indicated genotypes (n=30 per genotype). Note that *β*-lobe crossing characteristic of *Nab2* nulls (>80% penetrance, bar) is completely suppressed by loss of *Mettl3* ( bar).

### Nab2 binds *Sxl* pre-mRNA and modulates its m^6^A methylation

The ability of the *Mettl3^null^* allele to promote appropriate sex-specific exon 2-exon 4 splicing of *Sxl* in *Nab2^ex3^* females is significant, both because it identifies the m^6^A ‘writer’ Mettl3 as required for *Sxl* mis-splicing in heads lacking Nab2, and because this same *Mettl3* allele normally results in hypomethylation of *Sxl* mRNA and exon 3 inclusion in female flies (Haussmann et al., 2016; Kan et al., 2017; Lence et al., 2016). This paradox could be explained if exon 3 inclusion in *Nab2^ex3^* females is due to hypermethylation of the *Sxl* pre-mRNA, which is then suppressed by removing *Mettl3*. To test this hypothesis, a series of primer sets was designed to examine *Sxl* pre-mRNA and mRNA transcripts by RNA-immunoprecipitation (RIP) and anti-m^6^A-RIP (MeRIP) (**Figure 6A**). As illustrated in **Figure 6A**, the *Sxl* transcript contains candidate binding sites for both Sxl protein (polyuridine tracts=**red** ticks) and Nab2 protein (polyadenosine tracts=**green** ticks), and approximate sites of m^6^A methylation ticks) (mapped in Kan et al., 2017) (see **Supplemental Figure 7** for a complete schematic). To assess the m^6^A status of total *Sxl* RNA, MeRIP precipitates from female head lysates (control, *Nab2^ex3^*, *Mettl3^null^*, and *Nab2^ex3^*,*Mettl3^null^*) were analyzed by reverse transcription-real time quantitative PCR (RT-qPCR) with the exon 2-exon 2 (E2-E2) primer pair, which amplifies both pre-mRNA and mature mRNA (*Sxl^E2-E2^* in **Figure 6B**). This approach detects reduced *Sxl* m^6^A in *Mettl3^null^* heads relative to controls, which is consistent with prior studies (Haussmann et al., 2016; Kan et al., 2017; Lence et al., 2016), and an increase in *Sxl* transcript recovered from MeRIP of *Nab2^ex3^* heads, consistent with *Sxl* hypermethylation. This apparent increase in *Sxl* m^6^A methylation in *Nab2^ex3^* heads requires Mettl3, as *Sxl* mRNA recovery in MeRIP is strongly reduced in *Nab2^ex3^,Mettl3^null^* double mutant heads. Two m^6^A-methylated candidate Mettl3-target RNAs, *Act5c* and *Usp16* (Kan et al., 2017; Lence et al., 2016) were analyzed as additional positive controls for m^6^A status. MeRIP-qPCR indicates that both mRNAs are hypomethylated in *Mettl3^null^* and hypermethylated in *Nab2^ex3^* (**Supplemental Figure 8**). For *Act5c*, this *Nab2^ex3^* hypermethylation requires *Mettl3* (**Figure 6B**). Shifting this analysis to qPCR with the *Sxl* E2-E4 primer set (*Sxl^E2-E4^* in **Figure 6B**), which is predicted to selectively detect spliced *Sxl* mRNAs, supports a very similar pattern of elevated *Sxl* m^6^A in *Nab2^ex3^* heads that requires *Mettl3.* These MeRIP-qPCR data argue that Nab2 either inhibits Mettl3-mediated m^6^A deposition or promotes m^6^A removal on *Sxl* mRNAs, which in turn controls patterns of exon 3 retention/skipping. A prediction of this model is that Nab2 loss should result in hypermethylation of the *Sxl* pre-mRNA. Testing this hypothesis in MeRIP precipitates with the I3-E3 primer pair (*Sxl^I2-E3^* in **Figure 6C**) or the I3-E4 primer pair (*Sxl^I3-E4^* in **Figure 6C**) reveals moderate (1.5-fold) enrichment for intron 2-containing *Sxl* RNAs in *Nab2^ex3^* heads, and stronger (4.5-fold) enrichment for intron 3-containing RNAs, consistent with elevated m^6^A on *Sxl* pre-mRNAs that still contain introns 2 and 3. A parallel anti-Flag IP from head lysates of adult females expressing N-terminally tagged Nab2 specifically in neurons (*elav>Flag:Nab2*), coupled with RT-qPCR with I3-E4 primers, indicates that Nab2 associates with unspliced *Sxl* pre-mRNA (**Figure 6D**). In sum, these data provide a molecular framework to interpret *Nab2*-*Mettl3*-*Sxl* interactions in which Nab2 associates with the *Sxl* pre-mRNA, perhaps via the poly(A) sites located in I2 and I3 (**green** ticks; **Figure 6A**), and prevents its Mettl3-dependent hypermethylation, thus ensuring a level of m^6^A necessary to guide *Sxl* exon 3 in the developing female nervous system.

**Figure 6.**
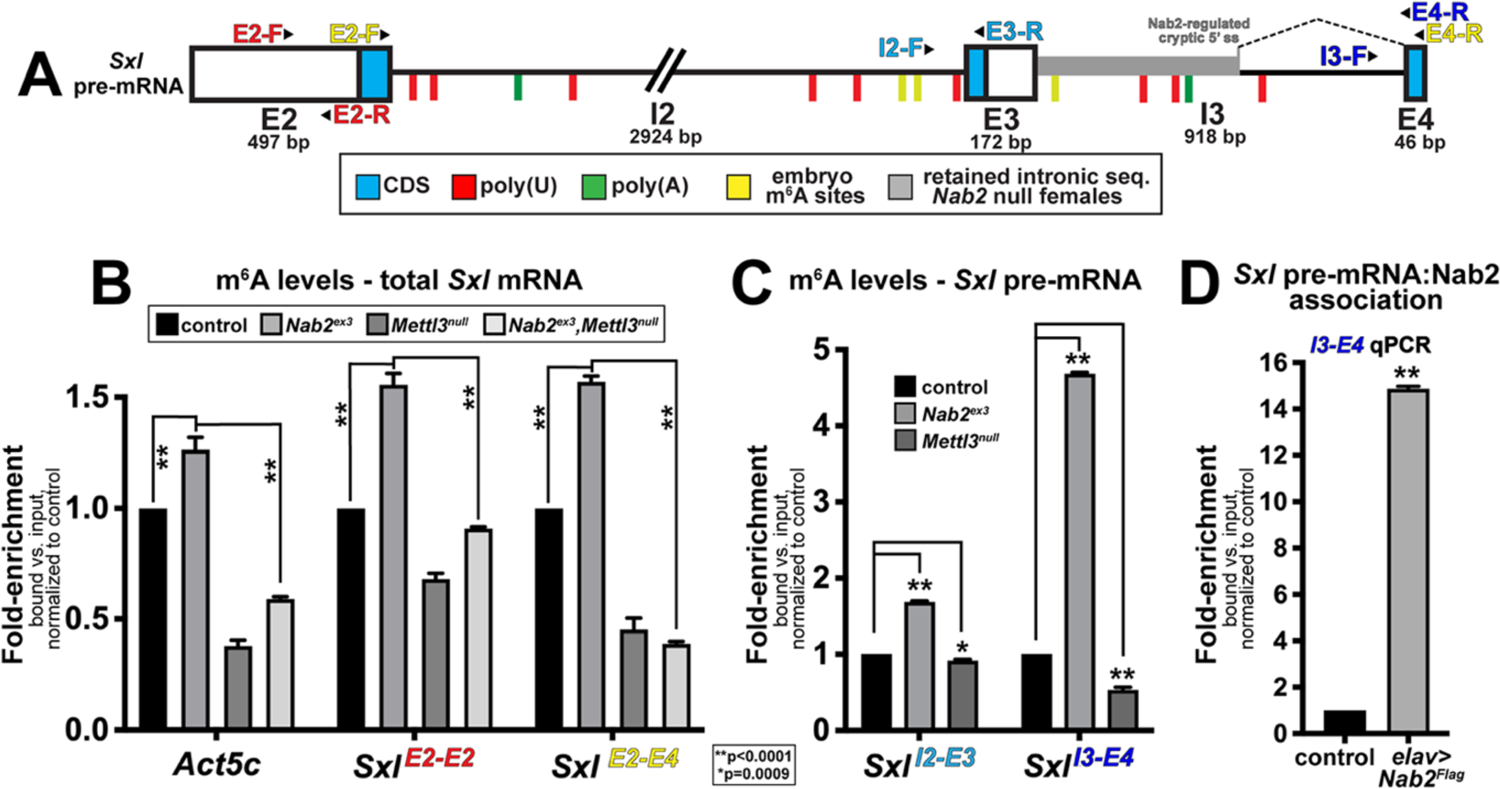
Nab2 associates with the *Sxl* mRNA and inhibits its m^6^A methylation. **(A)** Diagram of (E2, E3, E4) and introns (I2, I3) of the *Sxl* pre-mRNA annotated to show coding sequence (CDS; **blue**), the retained intronic region in *Nab2^ex3^* females (**grey**), and locations of color-coded primer pairs (**E2-F** and **E2-R**, and, **I2-F** and **E3-R**, **I3-F** and **E4-R**), poly(U) sites **red lines**, poly(A) sites **green lines**, and mapped m^6^A locations in *Drosophila* embryos (Kan et al., 2017). (**B)** Quantitative real-time PCR analysis of *Act5c* and *Sxl* mRNAs present in anti-m^6^A precipitates of control (*Nab2^pex41^*; black), *Nab2^ex3^* (**grey**), *Mettl3^null^* (**dark grey**), or *Nab2^ex3^*,*Mettl3^null^*) female heads. *Sxl* primer pairs are indicated (**E2-F**+**E2-R** and +). **(C)** Similar analysis as in **B** using **I2-F**+**E3-R** and **I3-F**+**E4-R** primer pairs to detect unspliced variants of *Sxl* mRNA in anti-m^6^A precipitates of control (black), *Nab2^ex3^* (**grey**) and *Mettl3^null^* (**dark grey**) female heads. **(D)** Quantitative real-time PCR analysis with the **I3-F**+**E4-R** primer pair in anti-Flag precipitates from control (*Nab2^pex41^*) and *elav-Gal4,UAS-Nab2:Flag* female heads. For all panels, 1-day old female heads were used in three biological replicates, and data represent bound vs. input ratios normalized to control (*Nab2^pex41^*). Significance values are indicated (\*\**p*<0.0001, \**p*=0.0009).

## Discussion

Here we have employed an unbiased high-throughput RNA sequencing approach to identify head-enriched RNAs whose levels or structure are significantly affected by Nab2 loss. Bioinformatic filtering of this high read-depth dataset reveals changes in levels and structure of a relatively small set of transcripts, with the latter effect on RNA structure traced to splicing defects - intron retention, alternative 5’ and 3’ splice site usage, and exon skipping - in a small group of approximately 150 mRNAs. Among these, the most significant change is female-specific inclusion of exon 3 in the *Sxl* mRNA and use of a cryptic 5’ splice site within the downstream intron. Because *Sxl* exon 3 contains a stop codon, its inclusion is predicted to disrupt female-specific expression of Sxl protein, a U-rich RNA binding protein that controls somatic and germline sexual identity via effects on splicing and translation of target mRNAs (rev. in Moschall et al., 2017; Penalva and Sanchez, 2003). Bioinformatic and genetic data indicate that *Sxl* mRNA may be an especially significant target of Nab2 in neurons: mis-spliced RNAs in *Nab2* mutant female heads are highly enriched for predicted Sxl binding motifs, and an allele of *Sxl* that constitutively skips exon 3 (Barbash and Cline, 1995) substantially reverses neurodevelopmental and behavioral defects in *Nab2* null females. Moving downstream of Sxl, alleles of male-specific dosage compensation complex (DCC) components, including the direct Sxl target *msl-2* (Bashaw and Baker, 1995, 1997), also rescue phenotypic defects in *Nab2* mutant females. Given that Msl-2 is not normally expressed or active in females, these data provide evidence that masculinized *Sxl* splicing and *msl-2*/DCC activation are important contributors to phenotypes in *Nab2* mutant female flies. These results imply a fairly specific link between Nab2 and the *Sxl* exon 3 splicing machinery, which is confirmed by strong genetic interactions between *Nab2* and the *Mettl3* methyltransferase that deposits m^6^A on *Sxl* pre-mRNA and promotes exon 3 skipping (Haussmann et al., 2016; Kan et al., 2017; Lence et al., 2016). Molecular assays provide key insight into these *Nab2*:*Sxl* interactions. The Nab2 protein associates with unspliced *Sxl* pre-mRNA in head lysates, and its loss results in Mettl3-dependent hypermethylation of mature *Sxl* mRNA and the unspliced *Sxl* pre-mRNA. Given the known role of m^6^A in regulating *Sxl* exon 3 splicing, these data imply that Nab2 interacts with the *Sxl* pre-mRNA in the nucleus and opposes excessive m^6^A methylation by the Mettl3 complex, thus ensuring a level of m^6^A necessary to guide *Sxl* exon 3 skipping in the developing female nervous system.

Our finding are consistent with a fairly specific role for nuclear Nab2 in control of exon-specific splicing patterns within a small subset of head RNAs, most of which are shared between males and females. One Nab2-regulated transcript, the sex determination factor *Sxl*, is a sex-specific target of Nab2 in the female nervous system, and evidence indicates that altered *Sxl* splicing contributes to contributes to their phenotypic defects. As Sxl is itself an RBP that can control splicing, some fraction of the mis-sliced mRNAs detected in our HTS approach may not be direct Nab2 targets, but rather Sxl targets. This hypothesis is supported by the enrichment for predicted Sxl-binding sites among mis-spliced mRNAs in *Nab2* mutant female heads, and by the substantial rescue conferred by the *Sxl^M8^* allele. However, splicing of the Sxl target *tra* is unaffected in the *Nab2* mutant RNA-Seq datasets. This could be due to lack of read depth, although this does not seem to be the case (see **Supplemental Figure 4**), or to Sxl-independent *tra* splicing in adult heads. Unbiased screens for Sxl target RNAs have carried out in ovaries (Primus et al., 2019) and primordial germ cells (Ota et al., 2017), but a similar approach has not been taken in the post-mitotic nervous system, where Sxl targets may differ from other tissue types. In this regard, the subgroup of Nab2-regulated head RNAs that also contain predicted U-rich Sxl binding motifs may be enriched for Sxl targets that contribute to developmental phenotypes in *Nab2* mutants.

The evidence for a *Drosophila* Nab2 role in splicing parallels evidence of accumulation of ∼100 intron-containing pre-mRNAs in *S. cerevisiae* lacking the *Nab2* homolog (Soucek et al., 2016), with only a few transcripts affected in yeast, similar to what is observed in the current *Drosophila* study. Rescue of *Nab2* mutants by neuron-restricted expression of human ZC3H14 (Kelly et al., 2014) implies that this specificity may be a conserved element of Nab2/ZC3H14 proteins in higher eukaryotes. Indeed, knockdown of ZC3H14 in cultured vertebrate cells results in pre-mRNA processing defects and intron-specific splicing defects in the few RNAs that have been examined (Morris and Corbett, 2018; Wigington et al., 2016). The basis for Nab2 specificity in *Drosophila* heads is not clear, but it could reflect a high degree of selectivity in binding to nuclear pre-mRNAs (e.g. *Sxl*), or to interactions between Nab2 and a second mechanism that defines splicing targets. The convergence of Mettl3 and Nab2 on the *Sxl* pre-mRNA represents the first evidence that Nab2 can modulate m^6^A-dependent control of pre-mRNA splicing. Hypermethylation of *Sxl* that results from Nab2 loss could reflect a requirement to bind A-rich sequences and block access of the Mettl3 complex or to recruitment of an m^6^A ‘eraser’. Intriguingly the human homolog of *Drosophila* Virilizer, a m^6^A methyltransferase subunit and splicing cofactor (Hilfiker et al., 1995; Niessen et al., 2001), was recovered in an IP/mass-spectrometry screen for nuclear interactors of ZC3H14 (Morris and Corbett, 2018), suggesting a potential functional interaction between these proteins on shared target RNAs. Additional evidence that Nab2 restricts m^6^A methylation of targets beyond *Sxl* (e.g. *Act5c* and *Usp16*; **Figs. 6B and S8**) highlights the possibility that Nab2 may have a broader role in modulating m^6^A-dependent RNA processing events, such as splicing, turnover, trafficking and translation.

The robust *Mettl3^null^* suppression of *Nab2^ex3^* mushroom body (MB) defects is unlikely to be mediated solely by effects on *Sxl*, which has no reported role in MB development. This observation suggests that Nab2 and Mettl3 have an overlapping set of target RNAs that extends beyond *Sxl*. As almost all *Nab2* mutant phenotypes originate from a Nab2 role in central nervous system neurons (Kelly et al., 2016; Kelly et al., 2014; Pak et al., 2011), suppression by *Mettl3^null^* also implies that the Nab2-Mettl3 functional interaction is likely to play out in neurons. The broad phenotypic rescue of *Nab2^ex3^* afforded by *Mettl3^null^* is consistent with a model in which loss of Nab2 leads to m^6^A hypermethylation of Nab2-Mettl3 shared targets, which is then suppressed by removing Mettl3. This ‘goldilocks’ model is consistent with the *Sxl* data presented here and implies that excessive m^6^A caused by loss of Nab2 perturbs neuronal mRNA processing and/or expression in a manner similar to m^6^A hypomethylation caused by loss of Mettl3. The precise effect of excess m^6^A is unknown, but it could result in excessive recruitment of m^6^A ‘reader’ proteins that overwhelm the specificity of downstream regulatory steps. Significantly, the ability of human *ZC3H14* to rescue *Nab2^ex3^* viability and locomotion when expressed in fly neurons indicates that this new m^6^A inhibitory role of Nab2 may be conserved within the Nab2/ZC3H14 family of RBPs, and that excessive m^6^A methylation of key RNAs also contributes to neurological deficits in *ZC3H14* mutant mice and patients.

## Materials and Methods

### *Drosophila* stocks and genetics

*Drosophila melanogaster* stocks and crosses were maintained in humidified incubators at 25°C with 12hr light-dark cycles. The alleles *Nab2^ex3^* (null), *Nab2^pex41^* (*precise excision 41*; control) and *UAS-Flag-Nab2* have been described previously (Kelly et al., 2014; Pak et al., 2011). Lines from Bloomington *Drosophila* Stock Center (BDSC): *GMR-Gal4 (*#1350), *elav^C155^-Gal4* (#458), *msl-2^227^* (#5871), *msl-2^kmA^* (#25158), *mle^9^* (#5873), *roX1^ex6^* (#43647). The *Mettl3^null^* allele was a kind gift of J-Y. Roignant. The *Nab2^ex3^*,*Mettl3^null^* chromosome was generated by meiotic recombination and confirmed by genomic PCR (Supplemental Figure 6).

### RNA Sequencing (RNA-Seq) on *Drosophila* heads

RNA-Seq was performed on three biological replicates of 60 newly-eclosed adult female and male *Drosophila* heads genotype (control and *Nab2^ex3^* mutants). Heads were collected on dry ice, lysed in TRIzol (ThermoFisher), phase-separated with chloroform, and ran through a RNeasy Mini Kit purification column (QIAGEN). Samples were treated with DNase I (QIAGEN) to remove DNA contamination and transported to the University of Georgia’s Genomics and Bioinformatics Core for sequencing. rRNA was depleted using a Ribo-Zero Gold Kit (Illumina) and cDNA libraries were prepared using a KAPA Stranded RNA-Seq Kit (Roche). Quality control steps included initial Qubit quantification along with RNA fragment size assessment on an Agilent 2100 Bioanalzyer before and after rRNA depletion. The cDNA libraries were then sequenced for 150 cycles on a NextSeq 500 High Output Flow Cell (Illumina) set to generate paired-end, 75 base-pair (bp) reads. Total sequencing yield across all samples was 81.48 Gbp, equivalent to about 1.1 billion reads in total and 91 million reads per sample. Sequencing accuracy was high; 93.52% of reported bases have a sequencing quality (Q) score greater than or equal to 30.

### Read mapping, differential expression, visualization of RNA-Seq dataset

Raw read FASTA files were analyzed on the Galaxy web platform (usegalaxy.org Afgan et al., 2018). The BDGP6 release *Drosophila melanogaster* genome (dos Santos et al., 2015) from release 92 of the Ensembl database (Yates et al., 2020) was used as input for subsequent read mapping, annotation, and visualization. Briefly, reads from all four NextSeq500 flow cell lanes were concatenated using the Galaxy *Concatenate datasets tail-to-head (cat)* tool and mapped using RNA STAR (Dobin et al., 2013) with default parameters with some modifications (available upon request). Mapped reads were assigned to exons and tallied using featureCounts (Liao et al., 2014) default parameters with some modifications (available upon request). Differential expression analysis was conducted for all 12 samples using DESeq2 (Love et al., 2014) (Galaxy Version 2.11.40.1) and default parameters with some modifications (available upon request). Differential exon usage was analyzed using Galaxy Version 1.20.1 of DEXSeq (Anders et al., 2012) and the associated Galaxy tool DEXSeq-Count in both “*prepare annotation*” and “*count reads*” modes. Both tools were run with the Ensembl GTF with default parameters with some modifications (available upon request). Unlike with DESeq2, female samples and male samples were compared in independent DEX-Seq analyses. Outputs from all of these tools were downloaded from Galaxy for local analysis, computation, and visualization.

For visualization, custom R scripts were written to generate volcano plots and heatmaps (available upon request). Additional R packages used include ggplot2 (Wickham, 2016) and ggrepel (Slowikowski, 2019). Scripts in the R programming language (Team, 2019) were written and compiled in RStudio (Team, 2018). Principal component analysis was conducted on Galaxy. Mapped reads were visualized in the Integrative Genomics Viewer (IGV) (Robinson et al., 2017) and annotated based on data available on Flybase (Thurmond et al., 2019). DESeq2 was used with default settings for transcript assembly to generate pie charts and correlation scatter plot. The resulting assembled transcripts were compared using DESeq2 to identify genes that change significantly (*p*-value<0.05, >1-fold change) in expression. Significant fold change values in either male or female from DESeq2 (adj. *p*-val<0.05 and |log_2_FC|>1) were plotted, with the color indicating the fold change threshold reached in either males or females. Significantly DE genes (adj. *p*-val<0.05 and |log_2_FC|>1) were classified by type, as indicated by their gene ID. Mixtures of isoforms (MISO) version 0.5.4 was run on aligned reads of length 76 bp, and then delta PSI values were computed for WT vs. mutant using miso for male and female samples separately with the flags --num-inc 10 --num-exc 10 --num-sum-inc-exc 50 --delta-psi 0.3 --bayes-factor 10.

### Mixtures of Isoforms (MISO) Analysis

MISO (Katz et al., 2010) was used to determine percent spliced in (PSI) values for annotated alternative 3’ splice sites, alternative 5’ splice sites, and retained introns for each sample separately as follows. Alternative splicing annotations were generated using the rnaseqlib (a direct link to script is listed here) (https://rnaseqlib.readthedocs.io/en/clip/) script, gff_make_annotation.py, with flags--flanking-rule commonshortest --genome-label dm6. Replicates for each sample were pooled, and only full-length, mapped reads (76 bp) were used for the MISO analysis since MISO requires all reads input to be of the same length. MISO was run with the flag –prefilter, and the output was then input into the script, summarize_miso, with the flag --summarize-samples. Next, differential, alternative 5’ and 3’ splice sites, and differential retained introns, were determined between *Nab2^ex3^* and control for males and females, separately, using the script, compare_miso, with flag --compare-samples. The output of compare_miso was then input into the script, filter_events, with the flags --filter --num-inc 10 --num-exc 10 --num-sum-inc-exc 50 --delta-psi 0.3 --bayes-factor 10, to obtain the final differential PSI values.

### Gene ontology (GO) analysis

Gene Set Enrichment Analysis (GSEA) software (Subramanian et al., 2005) was employed for Gene Ontology (GO) analysis. For clarity, analyses were conducted separately for each of the three top-level GO domains: *molecular function*, *biological process*, and *cellular component*. GSEA-compatible GO term gene sets for *Drosophila melanogaster* were acquired using the GO2MSIG web interface (Powell, 2014). GSEA Desktop for Windows, v4.0.3 (Broad Institute) was then used to identify two distinct classes of GO terms, independently for females and for males: (1) terms enriched among up- and downregulated transcripts in *Nab2^ex3^* compared to controls, and (2) terms enriched among transcripts alternatively spliced in *Nab2^ex3^* compared to controls. For the first class, inputs consisted of all genes whose expression could be compared by DESeq2 (i.e. adjusted *p*-value was not NA). For the second class, inputs consisted of all genes with known, annotated alternative splicing events according to MISO. To identify the first class of ontology terms, genes were ranked by log_2_ (fold change) calculated by DESeq2 and analyzed by the GSEA-Pre-ranked tool. To identify the second class of ontology terms, genes with were ranked by the absolute value of the difference in PSI (percent spliced in) between *Nab2^ex3^* and control calculated by MISO. This second ranking was analyzed by the GSEA-Preranked tool. Enriched GO terms (nominal *p*-value<0.05) identified for the first class were evaluated manually, surfacing multiple terms directly related to splicing. Enriched GO terms (nominal *p*-value<0.05) for the second class were ordered by normalized enrichment score (NES) and evaluated to identify the top “independent” GO terms. Terms were defined as “independent” by reference to their position in the GO hierarchy as reported on each term’s “Inferred Tree View” page of the AmiGO2 GO database web tool (Carbon et al., 2009). “Independent” terms had no parent, child, or sibling terms in the GO hierarchy associated with a higher NES than their own.

### RBPs Motif Enrichment Analysis using Mixture of Isoforms (MISO) Analysis

RNA sequences were taken at differentially retained introns and alternative 3’ and 5’ splice sites obtained from the MISO analysis on males and females separately (*Nab2^ex3^* mutants vs. control). The sequence for each of these went 25 bp into the exon(s) of interest and 1 kb into the intron of interest. In the case of alternative 3’ and 5’ splice sites, the sequences went 25 bp into the exon starting from the alternative spice site that is closest to the center of the exon (i.e. the inner-most splice site), and 1 kb into the intron starting from that inner-most spice site. To convert these to RNA sequences, DNA sequences were first obtained using fastaFromBed (Quinlan and Hall, 2010), and then all T’s were converted to U’s with a custom script. To obtain putative binding sites for RBPs at these sequences, they were then input into fimo using the flags --text --max-strand and the “Ray2013_rbp_Drosophila_melanogaster.meme” file (Grant et al., 2011).

### RNA isolation for reverse transcription (RT) PCR and real-time qPCR

Total RNA was isolated from adult tissues with TRIzol (Invitrogen) and treated with DNase I (Qiagen). For RT-PCR, cDNA was generated using SuperScript III First Strand cDNA Synthesis (Invitrogen) from 2μg of total RNA, and PCR products were resolved and imaged on 2% agarose gels (BioRad Image). Quantitative real-time PCR (qPCR) reactions were carried out in biological triplicate with QuantiTect SYBR Green Master Mix using an Applied Biosystems StepOne Plus real-time machine (ABI). Results were analyzed using the ΔΔCT method, normalized as indicated (e.g. to *Act5C*), and plotted as fold-change relative to control.

**Supplemental Table 1:**
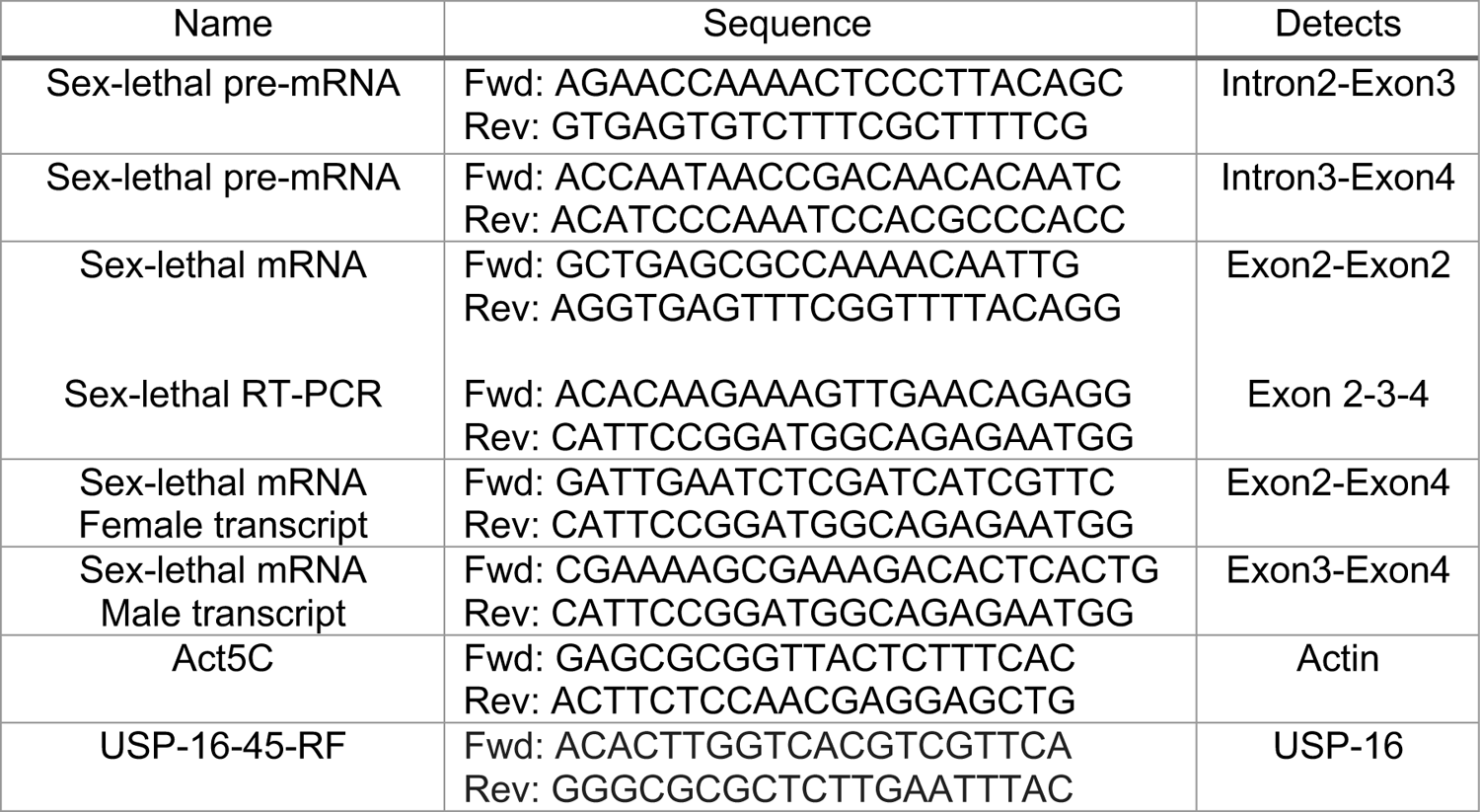
Primers used for RT and qPCR analysis

### Viability and lifespan analysis

Viability at 25°C was measured by assessing eclosion rates of among 100 wandering L3 larvae collected for each genotype and sex, and then reared in a single vial. Hatching was recorded for 5-6 days. At least 3 independents biological replicates per sex/genotype were tested and significance was calculated using grouped analysis on GraphPad (Prism). Lifespan was assessed at 25°C as described previously (Morton et al., 2020). In brief, newly eclosed animals were collected, separated by sex, placed in vials (10 per vial), and transferred to fresh vials weekly. Survivorship was scored daily. At least 3 independents biological replicates per vials of each genotype was tested and significance was calculated using grouped analysis on GraphPad (Prism).

### Locomotion assays

Negative geotaxis was tested as previously described (Morton et al., 2020). Briefly, newly eclosed flies (day 0) were collected, divided into groups of 10 male or females, and kept in separate vials for 2-5days. Cohorts of age-matched flies were then transferred to a 25-ml graduated cylinder for analysis. At least three biological replicates per sex were analyzed per genotype on GraphPad (Prism).

### Brain dissection and mushroom body (MB) imaging

Brain dissections were performed essentially as previously described in (Kelly et al., 2016). Briefly, adult brains were dissected in PBT (1xPBS, 0.1% TritonX-100) and collected in PBS at 4°C. Brains were then fixed in 4% paraformaldehyde at RT, washed 3x in PBS, and incubated in 0.3% PBS-T (1xPBS, 0.3% TritonX-100). Following blocking for 1hr (0.1% PBS-T, 5% normal goat serum), brains were stained overnight in block+primary antibodies. After 5x washes in PBT, brains were incubated in block for 1hr and transferred into in block+secondary antibody for 3hrs. Brains were then washed 5x in PBT and mounted in Vectashield (Vector Labs). The FasII antibody clone 1D4 (Developmental Studies Hybridoma Bank) was used to label MBs at a 1:50 dilution. Whole brain images were captured with a 20x objective. Maximum intensity projections were obtained by combining serial optical sections (Z-stacks) with Nikon A1R HD25 software using Fiji. The number of *α*-lobe and *β*-lobe defects (e.g. thin, missing or fused) were scored analyzed using GraphPad (Prism).

### Flag and m^6^A RNA immunoprecipitation (Flag-RIP and MeRIP)

The FLAG-RIP and MeRIP protocols were performed using previously described protocols in (Bienkowski et al., 2017) and (Lence et al., 2016) with some modification. Briefly, three replicates of 30 newly eclosed female flies were collected in 1.5 ml Eppendorf tubes and frozen in dry ice. Heads were removed with a 5.5 Dumont tweezer and homogenized with a mortar/pestle in Isolation buffer (50 mM Tris-HCl pH 8.1, 10 mM EDTA, 150 mM NaCl, and 1% SDS, 50 mM NaCl). This was diluted into IP buffer (50 mM HEPES, 150 mM NaCl, 5 mM EDTA, 0.5 mM DTT, 0.1% NP-40) supplemented with protease inhibitors (Roche) and RNasin Plus Inhibitor (Promega). Lysates were incubated with anti-Flag (M2 clone; Sigma) or anti-m^6^A (Synaptic Systems) antibodies and recovered on magnetic Protein G Dynabeads (Invitrogen). After overnight incubation at 4°C with rocking. Beads were washed 5x in IP buffer and RNA was isolated from antibody-bead precipitates, or controls (input samples) using TRIzol (ThermoFisher). Samples were treated with DNase-I and RNA was purified using RNeasy Kit (Qiagen).

### Statistical Analysis

Group analysis on biological triplicate experiments was done using Two-way ANOVA (Turkey’s multiple comparison test) on GraphPad (Prism) Version 8.4.2(464). Sample sizes (n) and *p*-values are denoted in the text or figures and noted by asterisks (for example, \**p*<0.05).

## Acknowledgments

Stocks obtained from the Bloomington Drosophila Stock Center/BDSC (NIH P40OD018537) were used in this study. We thank members of the Moberg and Corbett Labs for helpful discussions and advice, and B. Bixler, J. Tanquary, and C. Bowen for their contributions. We also thank M. Alabady (Georgia Genomics and Bioinfomatics Core) for technical support, and T. Lence and J.-Y. Roignant (Lausanne) for the gift of the *Mettl3* allele, and T. Cline (Berkeley) for discussions and insights on the *Sxl[M8]* allele. This work was funded by grants from the National Institute of Health to K.H.M. and A.H.C (R01 MH10730501), J.C.R. (F31 HD088043), and B.J. (F31 NS103595). B.J. and B.E.B were also supported during portions of the study by The Emory Initiative to Maximize Student Development (NIH R25 GM125598).

## Competing Interests

The authors declare no competing interests.

## Notes

https://www.ncbi.nlm.nih.gov/geo/query/acc.cgi?acc=GSE162531

### Competing Interest Statement

The authors have declared no competing interest.

### Summary of Updates

Added missing authors

