## Supplemental Figs and Legends for "The Nab2 RNA binding protein promotes sex-specific splicing of *Sex lethal* in *Drosophila* neuronal tissue"

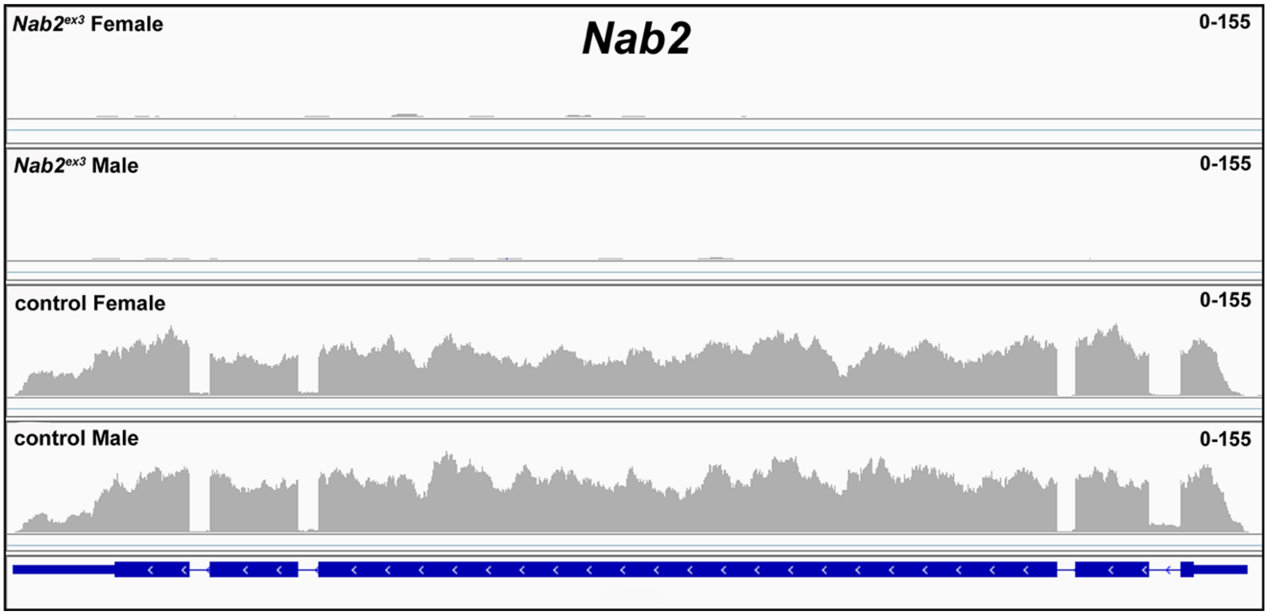


**Supplemental Figure 1. RNA sequencing reads across the *Nab2* locus**. IGV image of RNA sequencing reads across the *Nab2* locus in *Nab2*$^{ex3}$ (top tracks) and control (*Nab2*$^{pex41}$) adult female and male heads. Intron-exon structure is indicated at bottom. Read depth scale is indicated (0-155).

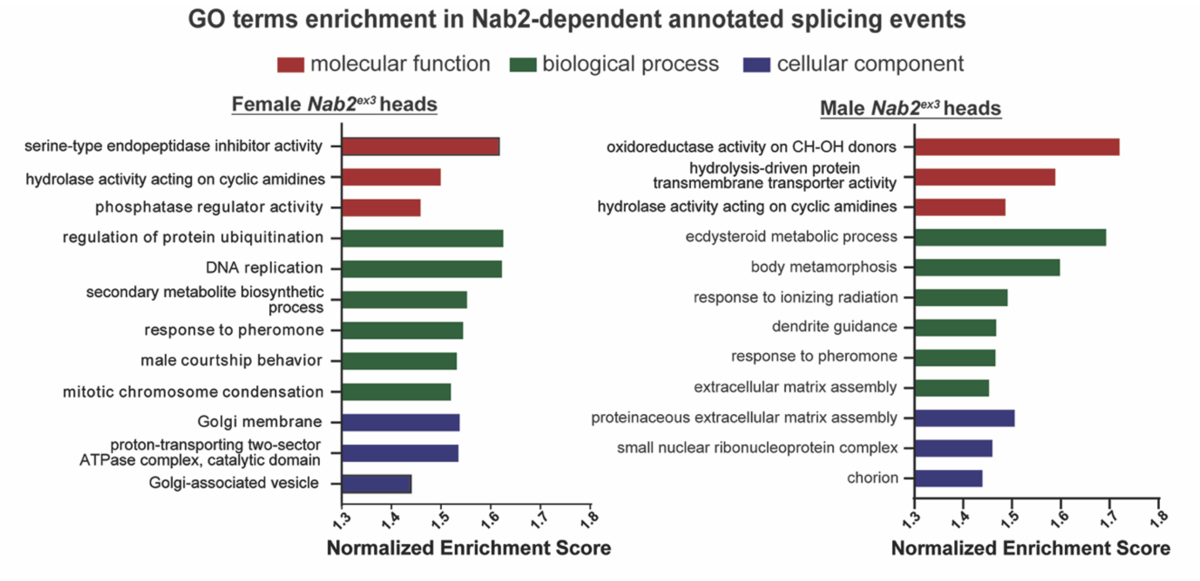


**Supplemental Figure 2. GO term enrichment among Nab2-regulated alternative splicing events**. Chart illustrating gene ontology (GO) terms enriched in the 'molecular function', 'biological process' and 'cellular component' categories among altered alternative splicing events detected by MISO in female and male *Nab2*$^{ex3}$ head RNAs.

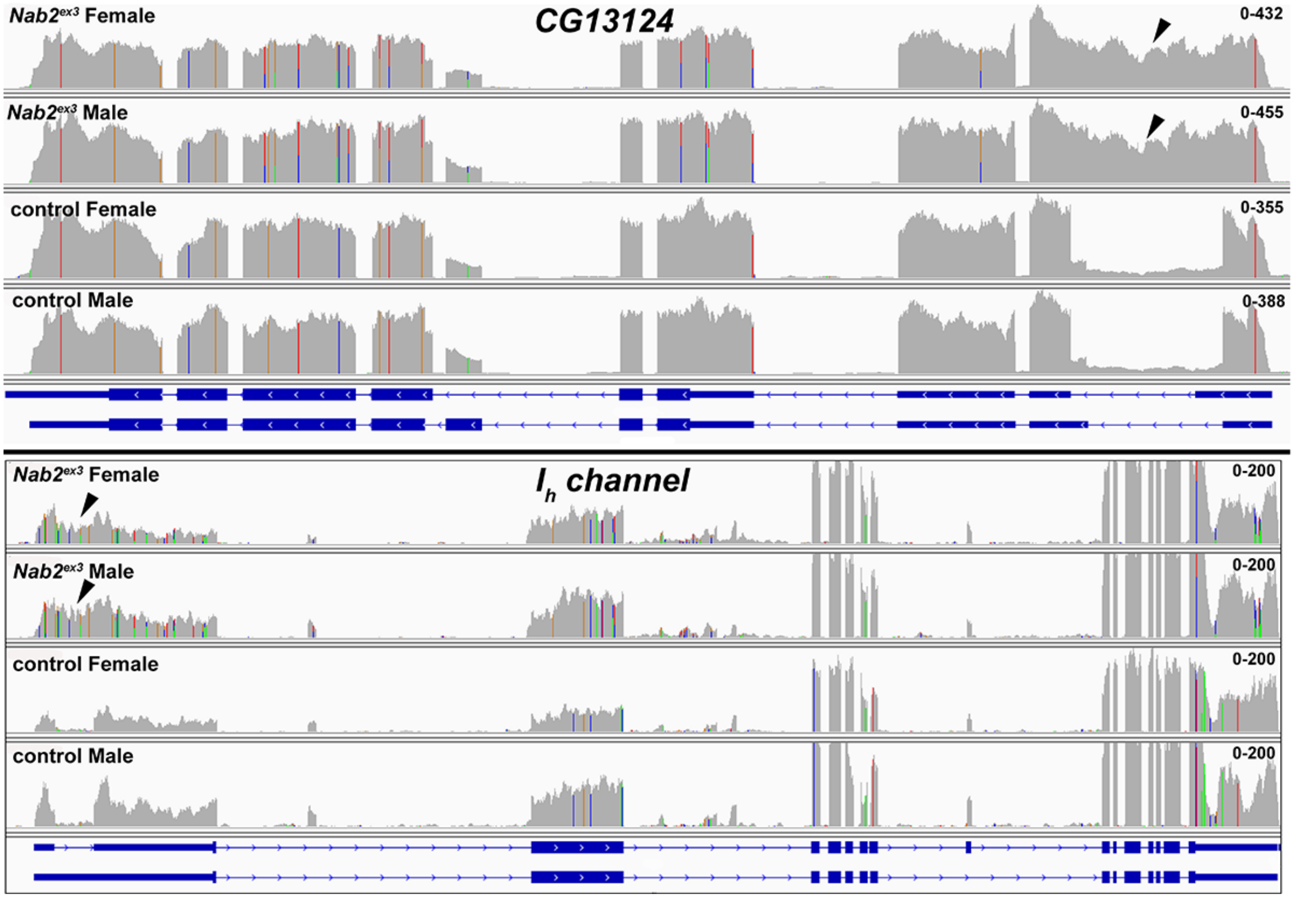


**Supplemental Figure 3. RNA sequencing reads across the *CG13124* and $I_h$ *channel* loci.** IGV images of RNA sequencing reads across *CG13124* and $I_h$ *channel* in *Nab2*$^{ex3}$ (top tracks) and control (*Nab2*$^{pex41}$) adult female and male heads. Intron-exon structure is indicated at bottom. Read depth scales are indicated. Arrowheads indicate reads across the first intron of each gene, consistent with intron-retention.

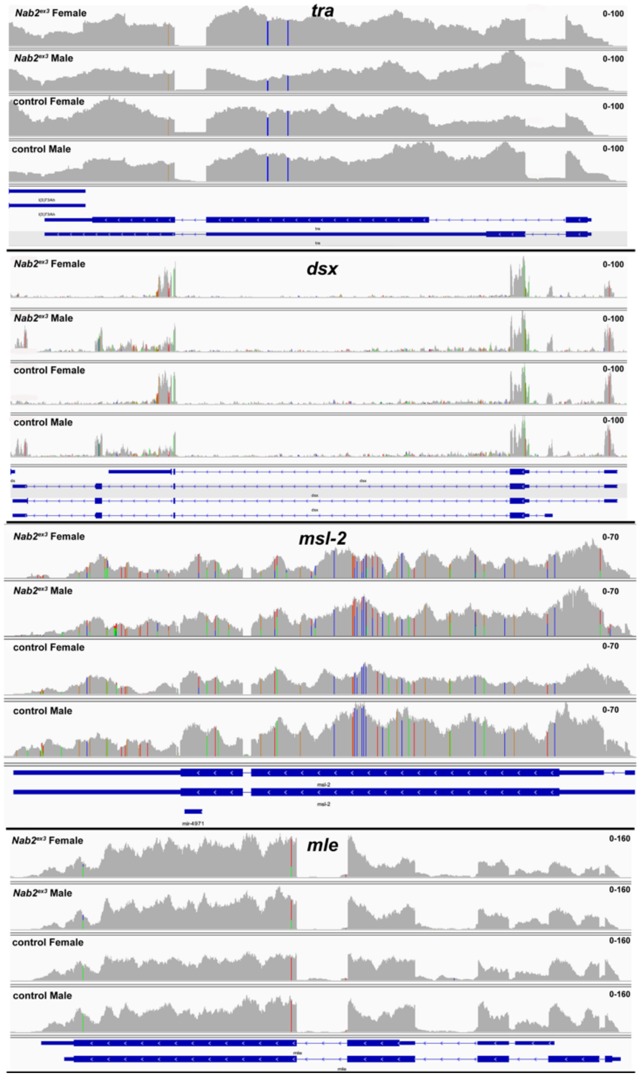


**Supplemental Figure 4. RNA sequencing reads across the *tra* and *dsx* loci.** IGV images of RNA sequencing reads across *tra*, *dsx*, *msl-2*, and *mle* in *Nab2*$^{ex3}$ (top tracks) and control (*Nab2*$^{pex41}$) adult female and male heads. Intron-exon structure is indicated at bottom. Read depth scales are indicated.

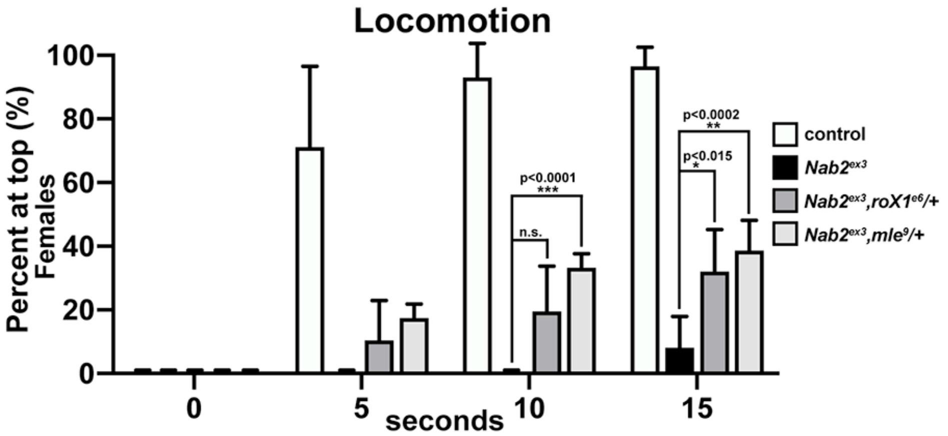


**Supplemental Figure 5. Modification of *Nab2$^{ex3}$* locomotor defect by *roX1* and *mle* alleles.** Negative geotaxis of age-matched adult female controls (*Nab2$^{pex41}$*), *Nab2$^{ex3}$* mutants, or *Nab2$^{ex3}$* mutants carrying single copies of the *roX1$^{e6}$* or *mle$^{9}$* loss-of-function alleles at 5sec, 10sec, and 15 sec timepoints. Significance values between indicated groups are indicated at the 30sec timepoint (p-values are indicated; n.s.=not significant).

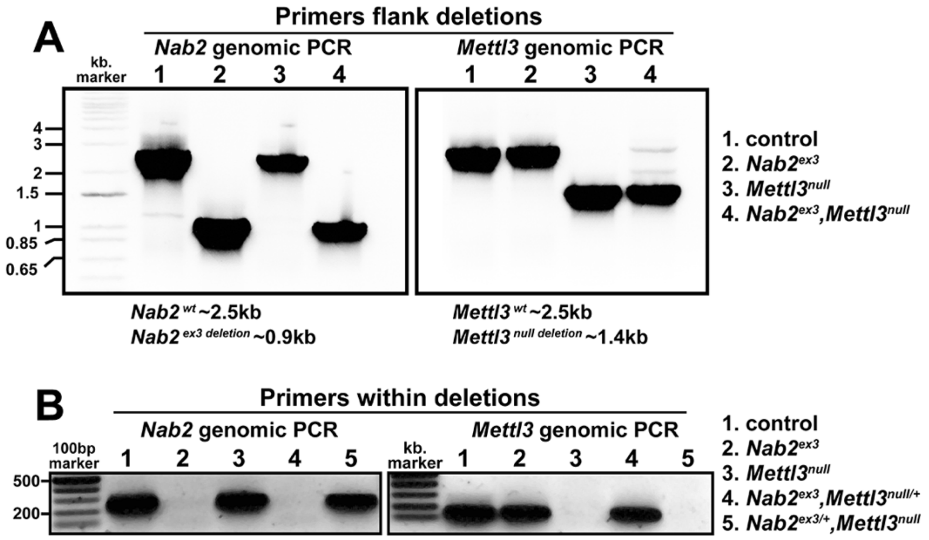


**Supplemental Figure 6. Genomic PCR confirms the *Nab2$^{ex3}$,Mettl3$^{null}$* recombinant**

Genomic PCR on the indicated genotypes using primer pairs that either flank **(A, top panel)** or lie within **(B, bottom panel)** the *ex3* deletion in *Nab2* (left half of each gel) or the *null* crispr deletion in *Mettl3* (right half of each gel). Approximate product sizes are indicated.

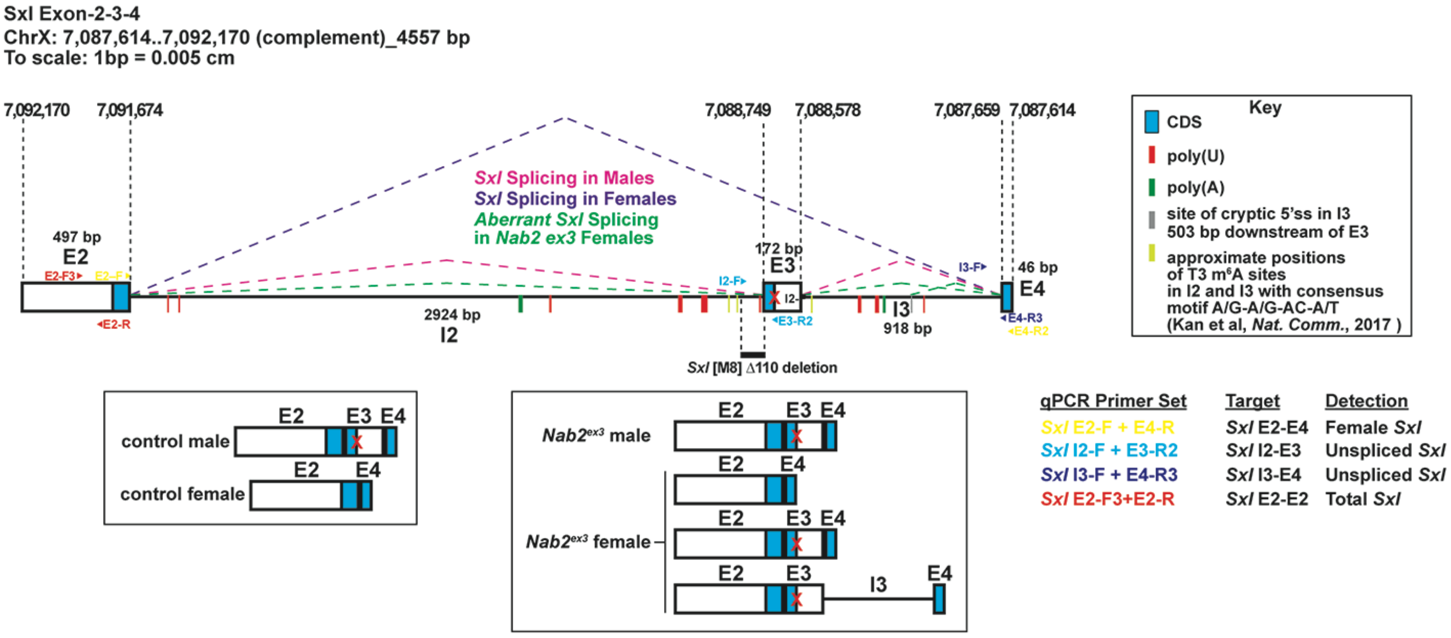


**Supplemental Figure 7. Genomic PCR confirms a *Nab2*$^{ex3}$,*Mettl3*$^{null}$ recombinant.** Detailed schematic of the exon 2-3-4 *Sxl* locus with annotated locations of introns and exons annotated to show coding sequence (CDS; blue), the retained intronic region in *Nab2*$^{ex3}$ females (grey), and locations of color-coded primer pairs (E2-F and E2-R, E2-F and E4-R, I2-F and E3-R, I3-F and E4-R), poly(U) sites red lines, poly(A) sites green lines, and mapped m$^6$A locations in *Drosophila* embryos yellow lines (Kan et al., 2017). Colored dotted lines indicated sex-specific splicing in wildtype adults and the altered splicing documented in this study. Boxed areas below summarize exon-intron structure in wild type heads and *Nab2*$^{ex3}$ heads. Base pair coordinates are indicated (Dm Release 6).

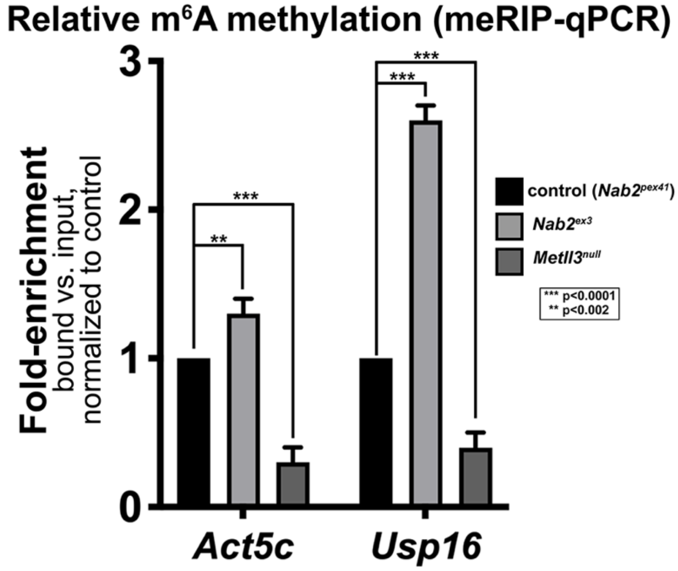


**Supplemental Figure 8. Nab2 limits m$^6$A methylation of additional Mettl3 target RNAs**

Quantitative real-time PCR analysis of *Act5c* and *Usp16* mRNAs present in anti-m$^6$A precipitates of control (*Nab2$^{pex41}$*; black), *Nab2$^{ex3}$* (grey), *Mettl3$^{null}$* (dark grey) adult female heads. 1-day old female heads were used in three biological replicates, and data represent bound vs. input ratios normalized to control (*Nab2$^{pex41}$*). p-values are indicated (n.s.=not significant).
